# Recurrent evolution of small body size and loss of the sword ornament in Northern Swordtail fish

**DOI:** 10.1101/2022.12.24.521833

**Authors:** Gabriel A. Preising, Theresa Gunn, John J. Baczenas, Alexa Pollock, Daniel L. Powell, Tristram O. Dodge, Jose Angel Machin Kairuz, Markita Savage, Yuan Lu, Meredith Fitschen-Brown, Molly Cummings, Sunishka Thakur, Michael Tobler, Oscar Ríos-Cardenas, Molly Morris, Molly Schumer

## Abstract

Across the tree of life, species have repeatedly evolved similar phenotypes. While well-studied for ecological traits, there is also evidence for convergent evolution of sexually selected traits. Swordtail fish (*Xiphophorus*) are a classic model system for studying sexual selection, and female *Xiphophorus* exhibit strong mate preferences for large male body size and a range of sexually dimorphic ornaments. However, sexually selected traits have been lost multiple times in the genus. Phylogenetic relationships between species in this group have historically been controversial, likely as a result of prevalent gene flow, resulting in uncertainty over the number of losses of ornamentation and large body size. Here, we use whole-genome sequencing approaches to re-examine phylogenetic relationships within a *Xiphophorus* clade that varies in the presence and absence of sexually selected traits. Using wild-caught individuals, we determine the phylogenetic placement of a small, unornamented species, *X. continens*, confirming an additional loss of ornamentation and large body size in the clade. With these revised phylogenetic relationships, we analyze evidence for coevolution between body size and other sexually selected traits using a phylogenetically independent contrasts approach. These results provide insights into the evolutionary pressures driving the recurrent loss of suites of sexually selected traits.

## Introduction

A fundamental puzzle in evolutionary biology is understanding the pressures that can lead to the recurrent evolution (or loss) of certain traits. Decades of work in evolutionary biology have studied convergent evolution in response to similar ecological pressures at the phenotypic (Reed *et al*., 2011), molecular (Nachman *et al*., 2003; Hoekstra *et al*., 2006; Zhen *et al*., 2012; Mohammadi *et al*., 2021), and genomic levels (Lee & Coop, 2019). Research has also highlighted how convergent evolution to similar ecological pressures can drive phenotypic shifts in several quantitative traits, resulting in distantly related species with shared suites of traits. Recent work in this area includes phenotypic shifts associated with pollination (Katzer *et al*., 2019; Wessinger *et al*., 2019), the evolution of Batesian mimicry (Kunte, 2009; Kunte *et al*., 2014; Nixon & Parzer, 2021), adaptation to similar ecological niches (Rennison *et al*., 2019), or even to similar social environments (Purcell *et al*., 2014). Understanding repeated shifts in phenotype in response to environmental pressures—especially when such shifts involve concurrent changes in several traits—is a key piece of the puzzle of how organisms adapt to their environments.

One area in which recurrent phenotypic evolution has not been well-studied is in the case of sexually selected traits. These traits are particularly interesting because they often experience conflicting selective pressures from sexual selection and natural selection. Theory posits that in species where males experience stronger sexual selection, large, ornamented males tend to be preferred by females and have higher fitness as a result of greater mating opportunities (Shuster & Wade, 2003; Møller, 2021; Rosenthal, 2017). However, the same ornaments that improve mating success can also reduce the probability of survival, often though an increase in the risk of predation (Hernandez-Jimenez & Rios-Cardenas, 2012; Okada *et al*., 2021); although other mechanisms exist (McKean & Nunney, 2008; McNamara *et al*., 2013; Moore *et al*., 2021).Variation in the relative costs and benefits of ornamentation can lead to a range of reproductive strategies over evolutionary timescales. For example, in groups of species where sexual selection is stronger on males, males of some species may evolve costly ornaments and others may lose ornamentation entirely. Ornamentation is frequently associated with behavioral traits such as courtship (Mitoyen *et al*., 2019), and in some taxa, lack of ornamentation is frequently associated with the use of coercive mating strategies (Ryan & Causey, 1989; Emlen, 1997; Taborsky, 1998; Abbott *et al*., 2019). These different mating strategies have evolved repeatedly (Zimmerer & Kallman, 1989; Gross, 1996; Neff, 2001; Corl *et al*., 2010), and often involve coordinated changes in suites of traits.

*Xiphophorus* are a classic model of sexual selection and an ideal system with which to study how suites of sexually selected traits evolve. Commonly referred to as swordtails, males in many species develop a long extension on their caudal fin referred to as the “sword” ornament (Darwin, 1871). The sword ornament is a composite trait: the fin extension is paired with one or two black stripes under independent genetic control (Powell *et al*., 2021), and in some species the sword itself is colorful (Kallman & Bao, 1987). The sword ornament is attractive to females, especially in combination with courtship displays (Rosenthal *et al*., 2001; Basolo & Trainor, 2002). Despite this, there have been several losses of the sword within the genus (Fig. 1). In some cases, these losses are associated with changes in mating behavior (Ryan & Causey, 1989; Morris *et al*., 2005), suggesting possible shifts in reproductive strategy.

**Figure 1.**
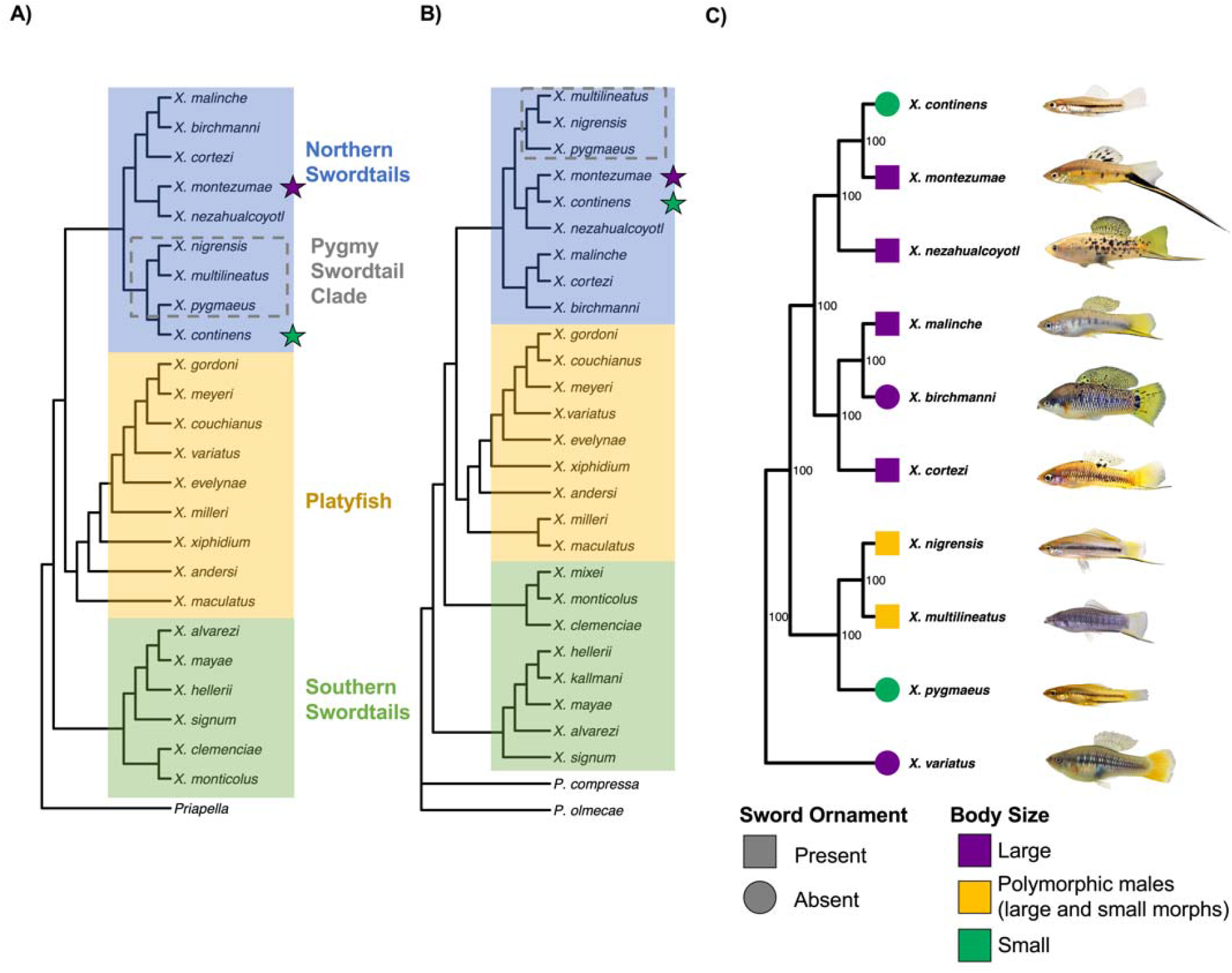
Phylogenetic relationships inferred in previous studies and in the present study. **A**) Summary of phylogenetic relationships inferred from previous genomic studies (green star highlights placement of *X. continens*, purple star highlights placement of *X. montezumae*). Whole genome phylogenetic analysis based on RNAseq and RADseq data (Cui *et al*, 2013 and Jones *et al*., 2013 respectively) placed *X. continens* sister to *X. pygmaeus*, within the “pygmy” swordtail clade. **B**) Earlier phylogenies using nuclear and mitochondrial markers and morphological characteristics placed *X. continen*s sister to *X. montezumae*. Shown here is a topology inferred with the dataset of Kang *et al*., 2013 (note: this tree was inferred using the authors’ alignment with RAxML instead of MEGAand jModeltest, which may account for slight differences in species placement). **C**) Phylogenetic relationships between Northern swordtail species inferred by this study using whole genome resequencing data for whole genome nuclear alignments. Analysis was performed using RAxML with the GTR+GAMMA model. Nodal support was estimated using 100 rapid bootstraps. Trees were rooted using the branch separating platyfish species from Northern swordtail species. Representative male phenotypes are shown next to the species names. For phylogenetic relationships inferred using mitochondrial sequences see Fig. S5.

The sword is not the only trait that is attractive to *Xiphophorus* females, and by examining coordinated evolution of other sexually selected traits, we can begin to disentangle how suites of sexually selected traits evolve. In *Xiphophorus* species tested to date (and in many other related species), females prefer to mate with larger bodied males [e.g. (Ryan & Wagner, 1987; Rosenthal & Evans, 1998; Cummings & Mollaghan, 2006; MacLaren & Rowland, 2006; Wong *et al*., 2011)]. Male *Xiphophorus* vary up to 3-fold in body size across species and some exhibit stable polymorphisms in body size within-species (Kallman, 1989; Ryan & Causey, 1989; Lampert *et al*., 2010). These stable male polymorphisms are known as male “morphs.” In both *X. multilineatus* and *X. nigrensis*, the larger morphs exhibit a suite of sexually selected traits, including the sword ornament, a deeper body shape, varied pigmentation patterns, and are more likely to exhibit courtship behaviors (Ryan & Causey, 1989; Zimmerer & Kallman, 1989; Liotta *et al*., 2019, 2020). Smaller morphs exhibit muted versions or the complete absence of these sexually selected traits and are more likely to engage in coercive mating tactics (often referred to as “sneaker” males). Similarly, in multiple species where males are fixed for especially small body size, there appears to have been concurrent loss or reduction of other sexually selected traits (Morris *et al*., 2005), but this hypothesis has not been systematically tested.

To formally test for convergent evolution of different sexually selected traits, we require an accurate phylogeny. The phylogeny of *Xiphophorus* has been the subject of frequent revision over the past several decades, and we revisit the phylogeny here focusing on species whose phylogenetic relationships may not be accurately resolved. Two species within one clade, the “Northern” swordtail clade, *X. pygmaeus* and *X. continens*, are the smallest and least ornamented species (Fig. 1). Two previous studies using whole-genome data placed these species as sister taxa (Fig. 1B; Cui *et al*., 2013; Jones *et al*., 2013). Earlier studies that used a variety of approaches including morphological traits, mitochondrial markers, and allozymes placed *X. continens* as the sister species of *X. montezumae* (Fig. 1A; Rauchenberger *et al*., 1990; Meyer *et al*., 1994; Morris *et al*., 2001; Kang *et al*., 2013). These findings were striking because *X. montezumae* is one of the most dramatically ornamented swordtail species (Fig. 1). The phylogenies of Cui *et al*. and Jones *et al*. (2013) directly contradicted this finding, including in analyses of mitochondrial markers, and the authors noted the discrepancy as a potential source of concern in the placement of *X. continens* (Cui *et al*., 2013). Interpretation of this result has been complicated by the fact that most studies have used long-maintained lab-stocks for *X. continens* (Table S1). While lab-stocks maintained by the *Xiphophorus* Genetic Stock Center and in individual research labs have been an invaluable resource for researchers for over 50 years, interfertility between nearly all species and high phenotypic similarity in some pairs of species raises challenges for maintaining long-term genetic resources.

In this study, we generate a phylogeny of *X. continens* and its relatives in the Northern swordtail clade, relying largely on whole-genome sequencing of wild-caught specimens. We find that *X. continens* is the sister species of the highly ornamented species *X. montezumae*, indicating a dramatic shift in reproductive strategy since the two species diverged (Fig. 1C). With the new phylogeny, we use phylogenetically independent contrasts to test hypotheses about the recurrent loss of suites of sexually selected traits across the genus.

## Methods

### Sampling and terminology

Throughout the manuscript, we refer to several different recognized clades of *Xiphophorus*, outlined in Figure 1. These include the Southern swordtail, Northern swordtail, and platyfish clades, which are the three major evolutionary lineages within *Xiphophorus* (Fig. 1). We also refer to the “pygmy” swordtail clade, which includes *X. pygmaeus* and its close relatives (Fig. 1).

Samples included in this phylogenetic analysis were a combination of previously published data from wild-caught Northern swordtail individuals and data generated for this project. The only species from which a wild caught sample was not available was the species *X. nigrensis* (see Supplementary Information 1; Fig. S1-S3). For this species we used a sample from the Brackenridge Field Laboratory at University of Texas at Austin where *X. nigrensis* individuals have been maintained in a colony derived from a wild-caught population since 2016. Sampling localities for the other specimens and data sources can be found in Table 1. All sequenced individuals were males.

**Table 1.** Sampling locations and data sources of genomic data for Northern swordtail species analyzed in this study.

| Species | Source population | Source data |
| --- | --- | --- |
| *X. malinche* | Chicayotla | Schumer et al. 2018 |
| *X. birchmanni* | Coacuilco | Schumer et al. 2018 |
| *X. cortezi* | Huichihuyán | Powell et al. 2021 |
| *X. montezumae* | Tamasopo | Schumer et al. 2016 |
| *X. nezahualcoyotl* | Los Gallitos | Schumer et al. 2016 |
| *X. nigrensis* | UT Austin – collection from Nacimiento de Río Choy | This study |
| *X. multilineatus* | Río Tambaque | This study |
| *X. pygmaeus* | Puente de Huichihuyán | This study |
| *X. continens* | Río Ojo Frío | This study |

### Whole genome resequencing

For this project, we generated data for four species where whole-genome data was not already available: *X. continens, X. pygmaeus, X. multilineatus*, and *X. nigrensis*, using a shearing-based library preparation protocol. DNA was extracted from fin clips using the Agencourt DNAdvance bead-based extraction protocol. The extraction method followed the manufacturer’s recommendations except that half-reactions were used. DNA was quantified using a Qubit fluorometer. The library preparation protocol used 500 ng – 1 ug of genomic DNA and followed the protocol developed by Quail et al. for Illumina library preparation (Quail *et al*., 2009). Genomic DNA was sheared to approximately 400 bp using a QSonica sonicator. Sheared DNA underwent end-repair via a 30-minute incubation at room temperature with dNTPs, T4 DNA polymerase, Klenow DNA polymerase and T4 PNK. Fragments were A-tailed with Klenow exonuclease and dATP via a 30-minute incubation at 37 °C, and adapters were ligated following this step. Purification was performed between each reaction step with a Qiagen QIAquick PCR purification kit. Unique barcodes were added to the libraries using indexed primers in a final PCR reaction using the Phusion PCR kit, with 12 cycles of amplification. This reaction was purified using 18% SPRI beads and resulting libraries were run on an Agilent 4200 Tapestation and quantified using a Qubit fluorometer. Libraries were sequenced on an Illumina HiSeq 4000 at Admera Health Services, South Plainfield, NJ. Raw sequence data has been deposited on the NCBI Sequence Read Archive (SRAXXXXX).

### Phenotyping

We were interested in generating a dataset where we could compare the evolution of sexually selected traits across *Xiphophorus*. Ideally, we would have access to photos of a number of individuals from wild populations. However, many species are only available as lab strains at the *Xiphophorus* genetic stock center, complicating interpretation of phenotypic variation within species. As a result, for our main analysis investigating coevolution between body size and male ornamentation we obtained photographs suitable for morphometric analysis from a single male and single female from each species and manually collected morphometric measurements in ImageJ 1.53k (Schneider *et al*., 2012) from each of them. For *X. multilineatus* and *X. nigrensis*, species which have large and small male morphs (Kallman, 1989), we collected data for one male of each morph. While some previous studies had suggested that other species might have more than one body size morph (Borowsky, 1987), we do not find clear support for this hypothesis in our analyses (see Supplementary Information 5). For each individual we measured standard length, sword length, dorsal fin length and height, peduncle depth, body depth, number of vertical bars, length and width of the lower and upper melanocyte pigmentation on the sword, peduncle pigmentation, as well as several binary traits (presence or absence of melanocyte pigmentation features, body color and sword color).

We were separately interested in quantifying phenotypic similarity between species that had similar ornamentation phenotypes but were distantly related based on our phylogenetic analysis (see Results). As a result, we generated an additional dataset for evaluating male phenotypic variation in *X. continens* and its relatives, as well as in *X. pygmaeus* and its relatives. For most of these species we were able to obtain morphometric photos of multiple wild-caught males (with the exception of *X. montezumae*, whose photos were provided by the *Xiphophorus* Genetic Stock Center). We measured the same phenotypes described above for a larger sample of *X. pygmaeus* (n = 11), *X. continens* (n = 6), *X. multilineatus* (n = 14; 7 per morph), *X. nigrensis* (n = 14; 7 per morph), and *X. montezumae* (n = 6).

*Xiphophorus* and related species have a reproductive strategy of internal fertilization and live birth. The gonopodium is a specialized male organ that is modified from the anal fin and is used to deposit sperm during internal fertilization. In addition to phenotypic measurements from pictures, we supplemented our dataset with measurements from the literature for gonopodial characteristics (Table S2; Jones *et al*., 2016; Supplementary Information 5). Several of these measurements were unavailable from the literature for the outgroup species (*Pseudoxiphophorus jonesii*). We also collected measurements from the literature, where available, on average male body size within *Xiphophorus* (Supplementary Information 5).

Sexual dimorphism is a frequently used proxy for the strength of sexual selection (Culumber & Tobler, 2017). To quantify sexual dimorphism, for each trait we took the difference between the male trait value and the female trait value in that species (following Culumber & Tobler, 2017). We normalized our measurements by standard length for traits that scale with body size (e.g. sword length, dorsal fin height). For example, in *X. malinche* the sword length divided by the body length was 0.13 in the focal male and 0 in the focal female, so the value for that trait for the dimorphism analysis in *X. malinche* was 0.13. We performed principal component analysis on a matrix of these male-female differences for all *Xiphophorus* species and an outgroup *(Pseudoxiphophorus jonesii)*.

### Variant calling and construction of alignments

To generate alignments for phylogenetic analysis, we first performed mapping of Illumina reads and variant calling. We mapped reads for each individual to the *X. birchmanni* reference genome (Powell *et al*., 2020a) using *bwa* (Li & Durbin, 2009). We identified and removed likely PCR duplicates using the program PicardTools (McKenna *et al*., 2010). We performed indel realignments and variant calling using GATK (version 3.4; McKenna *et al*., 2010) in the GVCF HaplotypeCaller mode. Past work has used mendelian errors in pedigrees to explore appropriate hard-call filtering parameters in *Xiphophorus* (Schumer *et al*., 2018). We used the thresholds identified by this work to filter variants based on a suite of summary statistics related to variant and invariant quality (DP, QD, MQ, FS, SOR, ReadPosRankSum, and MQRankSum; see Schumer *et al*., 2018). Based on the results of this previous analysis, we also masked all variants within 5 bp of an indel and all sites that exceeded 2X or were less than 0.5X the genome-wide background in coverage.

With this information in hand, we next turned to generating alignments for all northern swordtail species and outgroup species. Since all individuals were mapped to *X. birchmanni*, we generated pseudoreferences based on the *X. birchmanni* reference genome. Briefly, for each species, we used the *X. birchmanni* genome sequence and updated sites that were identified as variants and passed our quality thresholds, and masked all variant sites that did not pass our quality thresholds. We also masked invariant sites that failed quality thresholds that applied to both invariant and variant sites (for example, depth or proximity to INDEL filters). This resulted in pseudoreference sequences for 14 species (all Northern swordtails, two Southern swordtail species, and three platyfish species), aligned in the same coordinate space. Scripts and step-by-step examples for this workflow can be found online (scripts: https://github.com/schumerlab/Lab_shared_scripts; workflow: https://openwetware.org/wiki/Schumer_lab:_Commonly_used_workflows).

### Phylogenetic reconstruction with RAxML

For phylogenetic analysis, we extracted the twenty-four *Xiphophorus* chromosomes. We found that it was computationally intractable to analyze alignments with all sites included, so we took a two-pronged approach. For the analyses presented in the main text, we identified sites that were not monomorphic across the 14 focal species. We used these alignments as input into RAxML to build a total evidence phylogeny (Stamatakis, 2006). To construct the phylogeny, we conducted a rapid bootstrap analysis and searched for the best-scoring maximum-likelihood tree using a generalized time-reversible model (GTR+GAMMA) with 100 alternative runs on distinct starting trees. To perform analyses including invariant sites, we randomly sampled 1500 alignments 100 kb in length from the full dataset, representing approximately 20% of the genome. We analyzed this sub-sampled dataset as described above. These results mirror our results based only on variant sites (see Fig. S4). We separately analyzed an alignment of the mitochondrial genome (Fig. S5). We visualized results using the R packages ape, tidyverse, ggtree and associated packages (Yu *et al*., 2018; Paradis & Schliep, 2019; Wickham *et al*., 2019; Wang *et al*., 2020).

### Generating a merged phylogeny and performing phylogenetically independent contrasts

Using our newly developed phylogeny for Northern swordtails and previous phylogenetic results from our group for other *Xiphophorus* species (Cui *et al*., 2013), we generated a merged newick tree describing the inferred phylogenetic relationships between all *Xiphophorus* species, with the exception of *X. monticolus* and *X. kallmani*. Because our original phylogeny relied on coding sequences from RNAseq data (Cui *et al*., 2013) and our revised phylogeny for Northern swordtails is based on whole-genome sequences, we rescaled branch lengths to account for this (Supplementary Information 2). For the two species with multiple male morphs, *X. multilineatus* and *X. nigrensis*, we tried several different approaches and ultimately set the branch lengths within species to the length of the branch leading to the most recent common ancestor of *X. multilineatus* and *X. nigrensis* (Supplementary Information 2). This decision reduced possible issues identified in diagnostic tests that have been shown to lead to elevated false positive rates in phylogenetic analyses in previous work (Garland *et al*., 1992; Díaz-Uriarte & Garland, 1996); see Supplementary Information 2 for an in-depth discussion. We also used simulations to verify that the expected false positive rate in downstream phylogenetic analyses was not likely to be inflated (Supplementary Information 2).

Given the new placement of *X. continens* in the phylogeny (see Results), we were particularly interested in examining correlations between body size and other sexually selected traits. We leveraged our phylogeny and phenotypic data for each species to perform phylogenetically independent contrasts analysis for traits of interest (Felsenstein, 1985). Briefly, because a group of species shares a specific evolutionary history and hierarchy of relatedness, in some cases phenotypes are expected to be correlated simply as a result of their shared evolutionary histories. This non-independence generates statistical problems when testing for coevolution between traits of interest across species that vary in their relatedness to one another (Felsenstein, 1985). Phylogenetically independent contrasts methods attempt to correct for this non-independence using phylogenetic relationships between species and information about their divergence inferred from branch lengths (Garland *et al*., 1992).

For continuous traits, we used the R package *ape* to perform phylogenetically independent contrasts analysis. Before running these analyses, we first used the observed phylogenetic relationships between species, inferred branch lengths, and values for traits of interest, to perform diagnostic tests as proposed by Garland and colleagues (Garland *et al*., 1992); see Supplementary Information 3. We used the *ape* function *pic* to correct for phylogenetic signal in the traits of interest, including body size, sword index (calculated as sword length divided by standard length), dorsal fin index, sword edge width, number of vertical bars, and Principal Component 1 of sexual dimorphism (see Fig. 2). As recommended by theoretical work, we performed a regression through the origin to test for a correlation between the phylogenetically corrected traits of interest (Garland *et al*., 1992). We also used the R package *phytools* (Revell, 2012) to infer likely ancestral states for male body size, which was of particular interest for downstream analyses, using the *fastAnc* function.

**Figure 2.**
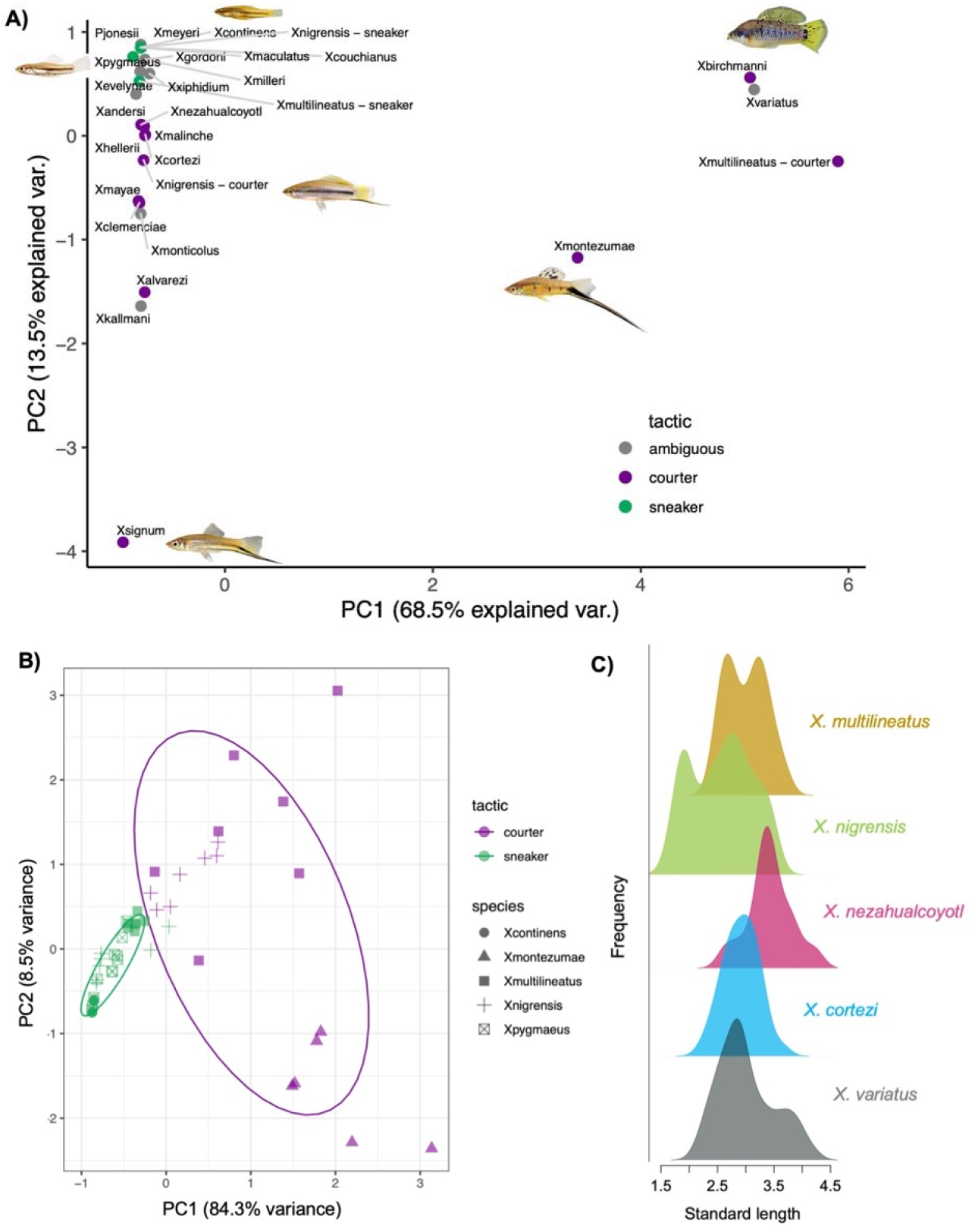
Sexual dimorphism across swordtails and analysis of male phenotypes and body size within Northern swordtails. **A**) Results of PCA analysis of sexual dimorphism among different *Xiphophorus* species and male morphs. See text for details on quantification of sexual dimorphism. **B)** PCA analysis indicates that *X. continens* and *X. pygmaeus* cluster with each other and with small morph males of *X. multilineatus* and *X. nigrensis*, but are phenotypically distinct from *X. montezumae* (the sister species of *X. continens*) and from large morphs of *X. multilineatus* and *X. nigrensis*. Color indicates mating strategy and shape indicates species. Ellipses indicate samples that falls within ± 1 standard deviation of the mean of that group. **C**) Standard length distribution of males from species that are known to have body size polymorphism (*X. nigrensis* and *X. multilineatus*), species that have not been reported to have body size polymorphism (*X. cortezi*), or species that have been previously suggested to have body size polymorphism (*X. variatus* and *X. nezahualcoyotl*). Color indicates species. The x-axis corresponds to standard length in centimeters. Sample sizes plotted per species are: *X. multilineatus* – n = 199, *X. nigrensis* – n = 60, *X. nezahualcoyotl* – n = 56, *X. cortezi* – n = 66, *X. variatus* – n = 56. A larger number of courter males compared to sneaker males was sampled for *X. nigrensis*; data was randomly downsampled for visualization.

We were interested in the relationships between body size and a subset of binary sexually dimorphic traits. In addition, certain traits in our dataset such as sword length take on continuous values but have a bimodal distribution (Fig. S6; see Supplementary Information 3). We analyzed several binary traits including the presence of the sword, the presence of the upper and lower sword edge, and a gonopodial trait (classification of ray 3 spine angle), using the method proposed by Ives and Garland (Ives & Garland, 2010) implemented in the R package *phylolm* (method=“logistic_IG10”; Tung Ho & Ané, 2014).

### Evaluation of demographic history, divergence and polymorphism in newly sequenced species

Several samples sequenced for this project represent wild-caught samples from species for which whole genome resequencing data has not been previously collected. For these species we calculated population genetic summary statistics including pairwise genetic divergence (D_xy_) between each species and their closest relative in our dataset and the θ _π_ estimate of genetic diversity within species.

We analyzed whole genome sequences using the pairwise sequentially Markovian coalescent (PSMC) approach (Li & Durbin, 2011) to infer changes in historical effective population size in *X. pygmaeus, X. continens*, and *X. multilineatus*. We excluded *X. nigrensis* from both this and the above analysis since wild caught samples were unavailable. In performing PSMC analysis, we assumed a generation time of 2 per year, a mutation rate of 3.5 × 10^−9^ per basepair per generation and a ratio of ρ/θ of 2, as we have for previous analyses of *Xiphophorus* demographic history (Schumer *et al*., 2018). Each species was analyzed separately. We performed bootstrap resampling of the data in bin sizes of 1 Mb to determine where we lose resolution to infer demographic history for each species in the recent and distant past.

### Analyses of gene flow

With newly available whole genome data, we were interested in re-examining patterns of gene flow within the Northern swordtail clade. Because we had aligned reads to a Northern swordtail assembly (*X. birchmanni*) for phylogenetic analysis, we were concerned about issues arising from reference bias that might generate similar signals to gene flow. To avoid this, we generated a new .vcf file using the *X. maculatus* genome (Schartl *et al*., 2013), which is an outgroup to all Northern swordtail species (Cui *et al*., 2013).

To do so, we indexed the *X. maculatus* genome using *samtools faidx* and *bwa index*. We then generated bam files using the same workflow and focal species as described above. Next, we performed variant calling for all of the *X. maculatus*-aligned bam files using *bcftools mpileup* to generate a joint .vcf file. We used vcftools to remove indels and sites with a minor allele frequency <5% (Danecek *et al*., 2011). The resulting .vcf was analyzed with the program *Dsuite* using the *Dtrios* and *Fbranch* commands to calculate Patterson’s D-statistic and F4 ratio statistics for each trio of Northern swordtail species (Malinsky *et al*., 2021), and distinguish between different branches as possible sources of phylogenetic discordance. For this analysis, we used *X. variatus* as an outgroup and provided Dsuite with the inferred genome-wide tree for Northern swordtails. We used a p-value threshold of 6×10^−5^ for the *Fbranch* analysis, corresponding to a Z-score of ∼4.

Dsuite analysis indicated strong evidence of gene flow between *X. continens* and an ornamented species, *X. nezahualcoyotl* (see Results). We were interested in polarizing the direction of this gene flow. However, existing approaches like *D*_*FOIL*_ (Pease & Hahn, 2015) could not be applied to this admixture event because of the branching order of the phylogeny. Instead, we took a different approach. We used PhyloNetHMM (Liu *et al*., 2014) to identify ancestry tracts that may have introgressed between *X. continens* and *X. nezahualcoyotl*, and examined patterns of divergence between pairs of species within these ancestry tracts to see if they were informative about the direction of gene flow. See Supplementary Information 4 for more details on this approach.

## Results

### An updated phylogeny for Northern swordtails

The total evidence phylogeny generated with RAxML had 100% bootstrap support at all nodes corresponding to species-level groups (Fig. 1C, Fig. S4; for mitochondrial results see Fig. S5). Most notably, our results using whole-genome sequencing data from a wild-caught *X. continens* sample place it sister to *X. montezumae* in the phylogeny. While this finding conflicts with previous results that used lab stocks (Cui *et al*., 2013; Jones *et al*., 2013), the phylogenetic placement of *X. continens* is concordant with older marker-based and morphological phylogenies, some of which used wild-caught *X. continens* samples (Table S1; Meyer *et al*., 1994; Morris *et al*., 2005; Kang et al. 2013). This indicates that the reproductive strategy of having unornamented, small males and using only coercive mating tactics (Ryan & Causey, 1989; Morris *et al*., 2005) evolved at least twice among Northern swordtails (Fig. 1C).

### Analysis of sexual dimorphism and traits correlated with body size in Xiphophorus

In contrast to previous findings (Cui *et al*., 2013; Jones *et al*., 2013), our revised phylogeny indicates that the evolution of both small body size and the loss of other sexually selected ornaments including the sword, vertical bars, and several pigmentation phenotypes, has occurred multiple times in *Xiphophorus*. With our revised phylogeny we infer three losses of the sword in Northern swordtails alone, and two instances of the evolution of extremely small body size (assuming that small body size arose once in the common ancestor of the pygmy swordtail clade and once in the lineage leading to *X. continens*; Fig. 1C). Ancestral state reconstruction analysis suggests that the ancestor of Northern swordtails was likely moderate in size, although confidence intervals for these estimates are large (Fig. S7).

Despite the phylogenetic placement of *X. continens* as sister species to *X. montezumae*, PCA analysis of phenotypic traits in *X. continens* and other swordtail species do not reflect this close relationship (Fig. 2A-2B). Instead, males of *X. continens* are grouped in PCA space with the small bodied males of other species, including *X. pygmaeus* and the small morphs of *X. nigrensis* and *X. multilineatus* (Fig. 2B).

To more formally explore which traits coevolve with body size in the revised *Xiphophorus* phylogeny, we used a phylogenetically independent contrasts approach (Felsenstein, 1985). We tested for correlations between phylogenetically corrected measures of body size and a number of traits that naively appear to correlate with body size in swordtails (i.e. without phylogenetic correction). Results for all traits analyzed are reported in Tables S3 & S4. We report p-values based on a Bonferroni correction for the number of tests in Tables S3 & S4 but discuss all results with uncorrected p-values <0.05 in the main text.

We first evaluated continuous traits. We found a strong relationship between body size and the number of vertical bars (Fig. 3A) and body size and dorsal fin index (Fig. 3C), traits that play a dual role in both mate choice and male-male competition. We also found a relationship between body size and the degree of sexual dimorphism within species based on PC1 of sexual dimorphism (Fig. 2A, Fig. 3B). We found a weaker but significant relationship between the sword length index and body size (Table S3). We also detected a significant relationship between sword edge pigmentation and body size (Table S3). Past research has highlighted the importance of the sword edge pigmentation in visual detection of the sword by females (Basolo & Trainor, 2002). Other continuous traits showed marginal or no significant associations with body size after phylogenetic correction (Table S3).

**Figure 3.**
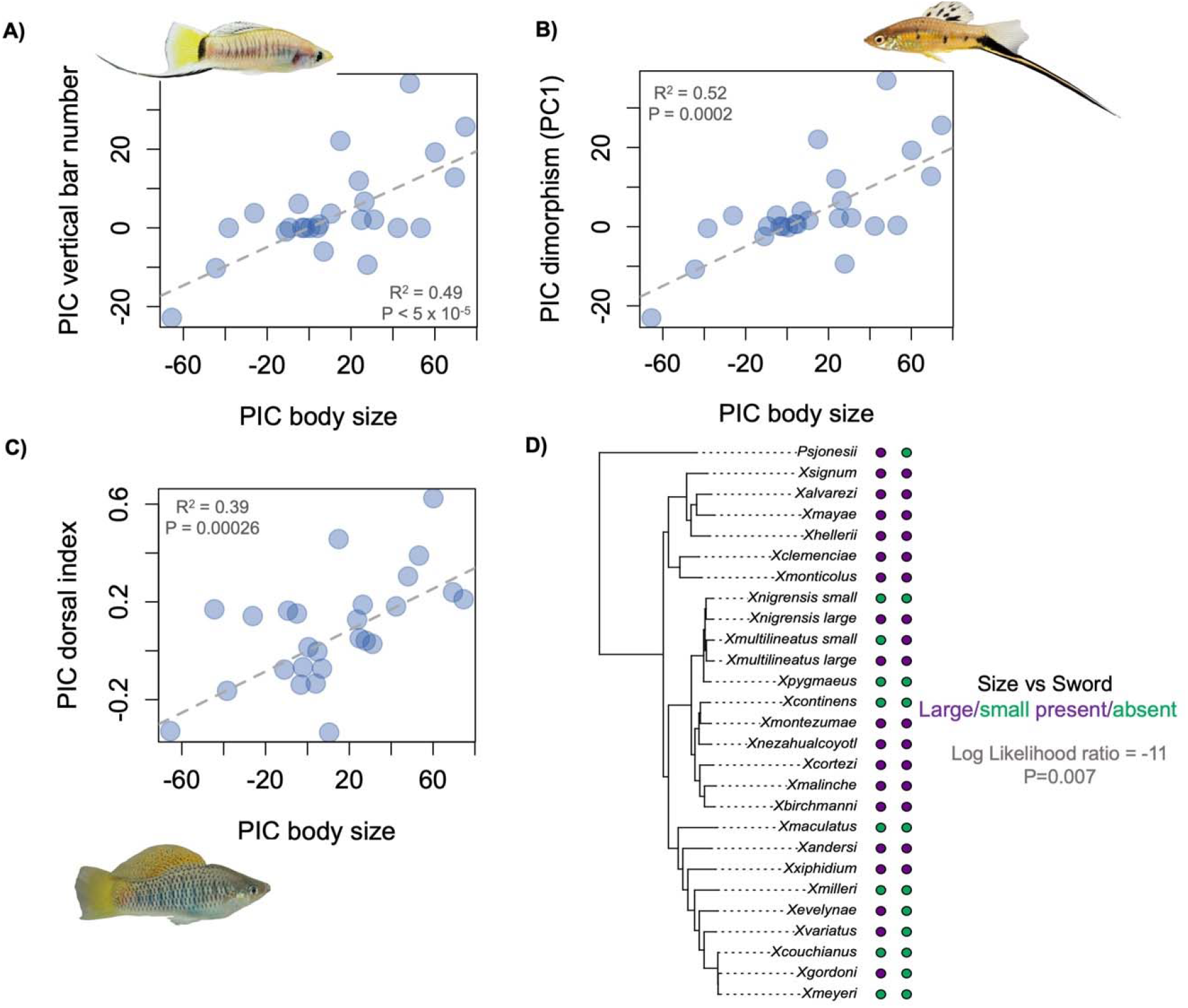
Results of phylogenetic independent contrasts analysis evaluating the correlations between body size and other sexually selected traits in the revised *Xiphophorus* phylogeny, after correcting for phylogenetic relationships between species. Results for all traits analyzed can be found in Tables S3-S4. **A**) We detect a significant correlation between body size and the number of vertical bars. A courter morph of *X. multilineatus* is shown in the inset. **B**) We also find a significant correlation between body size and PC1 of sexual dimorphism. A *X. montezumae* individual (with high sexual dimorphism), is shown in the inset. **C**) We detect a significant correlation between body size and dorsal fin index. *X. birchmanni*, a species with one of the highest scores for dorsal fin index, is shown. **D)** Using a phylogenetic logistic regression approach (Ives & Garland, 2010) we examined correlations between body size and binary traits. Shown here in colored dots next to the phylogeny is the relationship between body size (larger – purple, smaller – green) and sword presence (purple) or absence (green).

Analyzing the relationship between body size and binary traits, we observed a significant correlation between male size and the presence or absence of the sword (Log Likelihood ratio = - 11, p=0.0067; Fig. 3D), but not the presence or absence of the lower (Log Likelihood ratio= -13, p=0.082) or upper sword edge (Likelihood ratio=-13.4, p=0.052). Using a different threshold for sword presence or absence had some impact on the significance of our results but did not change qualitative patterns (Fig. S6; Supplementary Information 3).

We also analyzed gonopodial traits collected from the literature. While the function of variation in these traits is poorly understood, given their potential connection to mating strategy, we analyzed available data from the literature (Jones *et al*., 2016). We detected a significant relationship between the spine angle of ray 3 and body size (Table S4; Jones *et al*., 2016).

We repeated all analyses using body size measurements collected from the literature for a larger number of individuals and found that our results were generally concordant (Supplementary Information 5). We also performed analyses excluding either the small or large morphs of *X. multilineatus* and *X. nigrensis*. These results are reported in Supplementary Information 2 and Table S5.

### Population history of newly sequenced species

For our phylogenetic analysis we collected whole-genome resequencing data from wild-caught individuals of three species that had not been previously sequenced (apart from with RNAseq and reduced representation approaches; Cui *et al*., 2013; Jones *et al*., 2013): *X. pygmaeus, X. multilineatus*, and *X. continens*. Given the lack of previous data for these species, we report basic summary statistics on genetic diversity and divergence from their close relatives here and discuss inferences about their population history in more detail in the supplement (Supplementary Information 3; Fig. S8).

Like several other previously sequenced Northern swordtails, *X. continens* has very low genetic diversity, with a genome-wide θ_π_ estimate of 0.033% polymorphisms per basepair. This mirrors the low levels of genetic diversity previously reported in its closest relative, *X. montezumae* (θ_π_ = 0.03%; Schumer *et al*., 2016). Assuming that this level of diversity reflects the ancestral θ for the *X. montezumae* and *X. continens* clade (i.e. θ_A_ ∼ 0.03%), the estimated divergence time between *X. continens* and *X. montezumae* is 5.75 in units of 4*Ne* generations (D_xy_ = 0.38% per basepair). While *X. continens* and *X. montezumae* have similar levels of present-day nucleotide diversity, PSMC analysis suggests that *X. montezumae* had experienced a severe and sustained bottleneck over the last ∼10,000 generations (Fig. 4).

**Figure 4.**
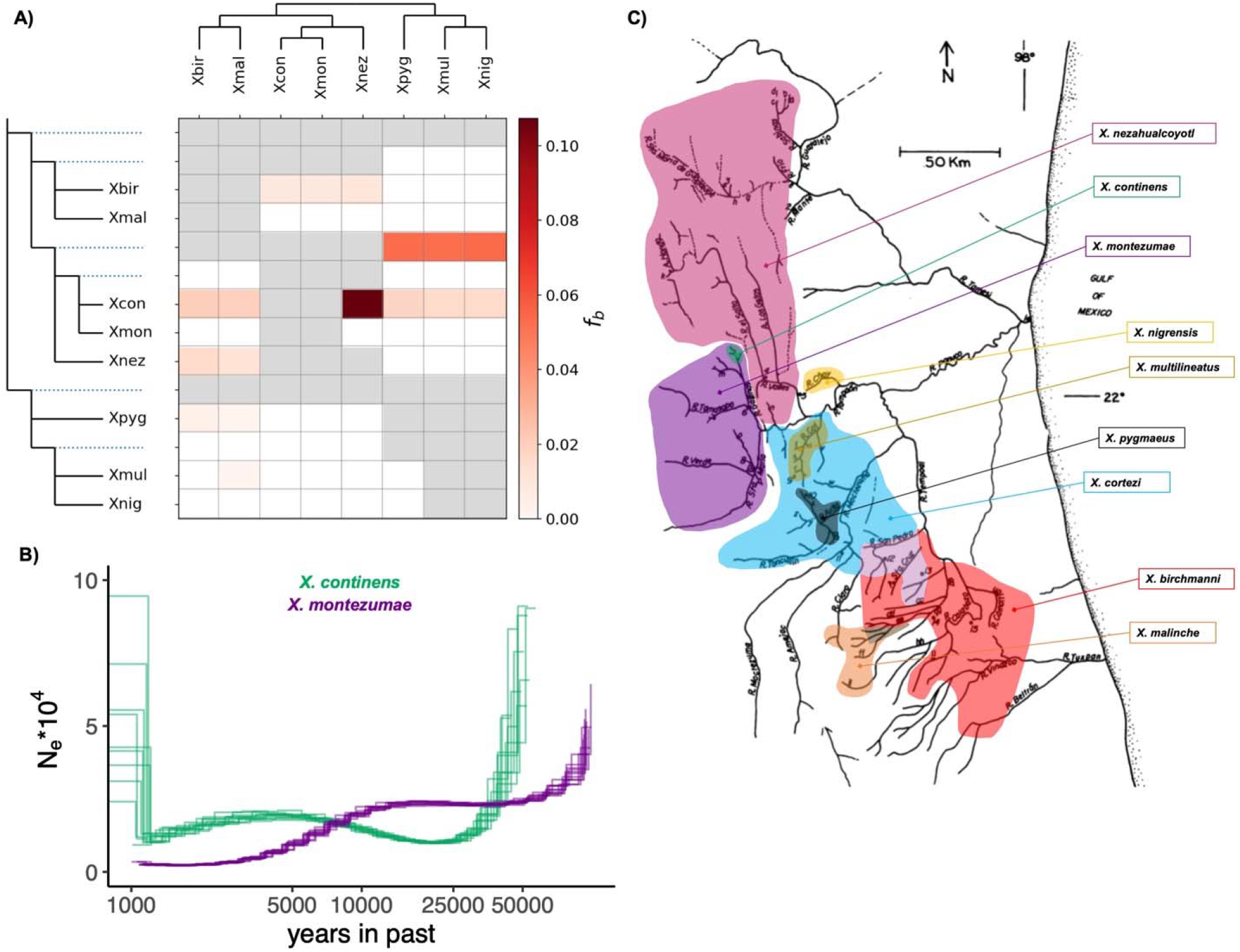
Gene flow estimates, species distributions, and demographic results for species of interest. **A**) Since Northern swordtail species are interfertile, we considered whether gene flow between the *X. pygmaeus* clade and *X. continens* could explain observed trait distributions. The gray squares indicate comparisons that could not be analyzed given the branching order of the phylogeny and the white squares indicate comparisons where no significant evidence of gene flow was found. Red hue squares indicate cases where evidence of gene flow is detected. The intensity of the red color corresponds to the value for the *f*_*branch*_ statistic calculated using *Dsuite*. The names of the focal species and their phylogenetic relationships are listed on the top and side of the matrix, blue dashed lines indicate comparisons involving the ancestral node. See Malinsky et al. for more information (Malinsky *et al*., 2021). These results show some evidence of admixture between the *X. pygmaeus* clade and *X. continens* but inferred levels of gene flow between these groups are low. Instead, *X. continens* is found to have substantial gene flow with *X. nezahualcoyotl*. For results including *X. cortezi*, which is inferred to have a history of gene flow with most species in the Northern swordtail clade see Fig. S9 and Supplementary Information 5. **B)** PSMC results estimating effective population size over time for sister species *X. continens* and *X. montezumae*. Analysis was conducted assuming a ρ/θ ratio of 2, generation time of two generations per year, and mutation rate of 3.5 × 10^−9^ following Schumer et al. 2018. Multiple lines reflect the results from 10 bootstrap replicates resampling 1 Mb segments. **C)** Range maps for Northern swordtail species; the original river map was adapted from Rauchenberger et al. 1990 and Cui et al. 2013.

*X. multilineatus* and *X. pygmaeus* have substantially higher levels of genetic diversity (θ_π_ of 0.071% and 0.073% respectively). Assuming that the ancestral θ was similar to present day θ in in this clade (θ_A_ ∼ 0.07%), the estimated divergence time between *X. multilineatus* and *X. pygmaeus* is approximately 1.9 in units of 4*Ne* generations (D_xy_ = 0.33% per basepair). The periods of inferred population growth and contraction in these two species differ (Fig. S8) and are discussed in more detail in Supplementary Information 3.

### History of admixture

The phylogenetic placement of *X. continens* indicates that this species either independently lost large male size, courtship, and male ornamentation traits, or that loci responsible for the loss of these traits spread from other species. To investigate this possibility, we used the program Dsuite to scan for evidence of gene flow across Northern swordtails (Malinsky *et al*., 2021). We calculated D-statistics for each triplet of species and used the version of the F4-ratio test implemented through the *Fbranch* command in Dsuite to explore admixture proportions and likely sources of gene flow within the Northern swordtail clade. These analyses confirmed several previously reported patterns of gene flow between species (Fig. 4; Supplementary Information 4; Cui *et al*., 2013; Schumer *et al*., 2016).

Surprisingly, we found very little signal of gene flow between *X. continens* and other Northern swordtails with small male body size (Fig. 4A; *X. pygmaeus, X. multilineatus*, and *X. nigrensis*) and instead found substantial evidence of gene flow with *X. nezahualcoyotl*. While this genetic exchange makes sense given their geographic proximity (Fig. 4C), *X. nezahualcoyotl* is an ornamented species with larger body size and lacks multiple male morphs (Fig. 2C).

Although our results are not suggestive of gene flow driving the transfer of alleles related to smaller body size and a lack of ornamentation, we note that these genome-wide test do not completely rule out the hypothesis. We detect low levels of gene flow between *X. continens* and species with small male morphs (Fig. 4A). Future studies could test this hypothesis more rigorously by constructing local phylogenies around the genes that underlie traits of interest once their genetic architecture is better understood.

Our analyses of putatively introgressed ancestry tracts suggested that the direction of gene flow was likely from *X. continens* into *X. nezahualcoyotl*. This pattern is notable because we previously found that *X. nezahualcoyotl* has substantial genetic contribution from *X. cortezi* as well (Schumer *et al*., 2016), implicating complex admixture in the evolutionary history of *X. nezahualcoyotl*. We discuss this result in more detail in Supplementary Information 4. More generally, *X. cortezi* is inferred to have extensive gene flow with many other species in the Northern swordtail clade, consistent with its widespread distribution (Fig. 4C). Given the complexity of gene flow involving *X. cortezi*, Fig. 4 shows the results without *X. cortezi* but we present the results of admixture involving this species in Fig. S9 and in Supplementary Information 4.

## Discussion

In our revised phylogeny of Northern swordtails, we find that *X. continens*, a small, unornamented species, is the sister lineage of *X. montezumae*, among the most ornamented species in the genus (Fig. 1). This means that *X. continens* has evolved small body size since it diverged from its common ancestor with *X. montezumae* (an estimated 450 thousand generations ago). The evolution of small body size in *X. continens* is accompanied by the loss of all other ornaments found in its close relatives, including the iconic sword ornament. The loss of the sword ornament in the *X. continens* lineage represents the fourth time across the entire *Xiphophorus* phylogeny that the sword and corresponding phenotypes (e.g. sword edge pigmentation) have been lost, and the third time this has occurred within the Northern swordtail clade. Using a phylogenetically independent contrasts approach, we infer that the patterns found in *X. continens* are generalizable across *Xiphophorus*: the evolution of smaller body size coincides with the loss or reduction of suites of other sexually selected traits. In addition to the loss of the sword and sword pigmentation, we find that the evolution of smaller body size coincides with a reduction in vertical bars, dorsal fin size, changes in gonopodial morphology, and a reduction in overall levels of sexual dimorphism (Table S3-S4).

Our whole-genome dataset allows us to address certain hypotheses about the genetic mechanisms underlying the recurrent evolution of small, unornamented males in *Xiphophorus*. Introgression has been shown to underlie patterns of recurrent phenotypic evolution in other species groups (Heliconius Genome, 2012; Jones *et al*., 2018; Oziolor *et al*., 2019), and past work has underscored the frequency of hybridization in *Xiphophorus* (Cui *et al*., 2013). We reexamined patterns of gene flow in Northern swordtails using our whole-genome dataset, and recapitulate several patterns of genetic exchange found by previous studies (Cui *et al*., 2013; Schumer *et al*., 2016). We see little evidence of genetic exchange between pairs of unornamented species such as *X. pygmaeus* and *X. continens*. This is despite the fact that *X. continens* and *X. pygmaeus* are strikingly similar at the phenotypic level (Fig. 2)—so similar as to have caused species misidentification in lab strains. Indeed, the only species inferred to have high levels of gene flow with *X. continens* is *X. nezahualcoyotl*, whose range is adjacent to that of *X. continens* (Fig. 4). This suggests that gene flow does not underlie recurrent loss of ornamentation in Northern swordtails, although future work should test this hypothesis at individual loci underlying particular traits (e.g. those associated with male body size; see Lampert *et al*., 2010).

The results of our demographic analysis are also not consistent with the hypothesis that species like *X. continens* and *X. pygmaeus* might have lost ornamentation traits due to genetic drift in small populations (see Supplementary Information 4 for in-depth discussion). The case of *X. pygmaeus* is especially interesting since its relatives have maintained a polymorphism for large, ornamented males with courtship behavior. This includes *X. multilineatus*, whose range is adjacent to that of *X. pygmaeus* (Fig. 4). Large historical effective population sizes in *X. pygmaeus* (Fig. S8) are instead suggestive of changes in the costs or benefits courtship and ornamentation in the ancestors of *X. pygmaeus*.

Classic research in sexual selection has underscored the importance of trade-offs between traits that facilitate survival and those that facilitate reproduction. Elaborate ornaments can increase an individual’s mating success while simultaneously reducing their probability of survival. The best studied sexually selected trait in *Xiphophorus* is the sword ornament (Darwin, 1871). Studies have indicated that females of most *Xiphophorus* species and those of related species strongly prefer the sword ornament, likely increasing the reproductive success of males with the trait (Morris *et al*., 1995; Basolo and Trainor 2001). However, the benefits of ornaments for mating success are accompanied by costs for survival, since it has been shown experimentally to attract predators and reduce critical swimming speed (Rosenthal *et al*., 2001; Kruesi & Alcaraz, 2007). In particular, *X. montezumae* has the longest sword ornament of all *Xiphophorus* species and our PCA analysis indicates that it is among the most sexually dimorphic species in the genus (Fig. 2; Kruesi & Alcaraz, 2007). Changes in the relative costs and benefits of ornamentation–for example, shifts in the ecological environment–could also explain the repeated evolution of small, swordless males (and the coincident loss of other sexually dimorphic traits; Fig. 3). Little is known about the ecological environments in which the least ornamented Northern swordtail species, *X. continens* and *X. pygmaeus*, are found, but anecdotal accounts suggest that they may live in faster flowing waters than many of their relatives (Rauchenberger *et al*., 1990).

Beyond the sword, we also infer a strong correlation on a phylogenetic scale between body size and dorsal index (Fig. 3C) and body size and male vertical bar number (Fig. 3A). Larger dorsal fins relative to male body size have evolved in several *Xiphophorus* species and are especially pronounced in *X. birchmanni* (Fisher *et al*., 2009). Females of some species appear to prefer large dorsal fins, perhaps because it contributes to larger perceived male size (MacLaren & Daniska, 2008; MacLaren *et al*., 2011). There is mixed evidence for direct preference for the dorsal fin itself (Robinson *et al*., 2011; Culumber & Rosenthal, 2013). In some species the dorsal fin is also important in male-male aggressive displays (Fisher & Rosenthal, 2007), so coevolution with large male body size could be driven by female preferences or by male-male competition. Vertical bars are a sexually dimorphic pigmentation pattern found in male *Xiphophorus*. These pigmentation patterns are multifunctional: they can deter aggression from conspecific males and simultaneously attract females (Morris et al. 1995). Within species with multiple male morphs like *X. multilineatus*, the number of vertical bars is more strongly predictive of male body size than other sexually dimorphic traits (Zimmerer & Kallman, 1989). Males darken vertical bars while engaging in courtship (Morris *et al*., 1995, 2008), directly linking this trait to a courtship strategy. In *X. multilineatus*, the absence of vertical bars is associated with small morph males that tend to exhibit coercive mating strategies (Morris *et al*., 2008). Like the sword ornament, large dorsal fins and vertical bars are thought to make males more conspicuous, and may similarly increase the risks of attracting predators.

It is interesting to speculate about connections between the evolution of these morphological phenotypes and the behavioral phenotypes observed in *X. continens* and in small males in the pygmy swordtail clade. Presumably once coercive mating strategies have arisen, the benefits of maintaining ornaments in individuals with this mating strategy are dramatically reduced while the costs remain, potentially explaining the coordinated loss of these suites of traits. Moreover, female preferences can also evolve, providing another possible mechanism that could drive the loss of ornamentation. For example, changes in female preference are thought to be responsible for the loss of the sword in the *X. birchmanni* lineage (Wong and Rosenthal 2006). However, changes in female preference do not provide a clear explanation for the evolution of small body size and the loss of ornaments in *X. pygmaeus* and *X. continens*. In *X. pygmaeus*, females prefer ornamented heterospecific males over unornamented conspecifics (Ryan & Wagner, 1987), although they discriminate against heterospecific males with vertical bars (Hankison & Morris, 2002, 2003), and females of some populations retain preferences for large male body size (Morris *et al*., 1996). In *X. continens*, females do not retain preferences for large male body size but show variation in preference for other ornaments found only in heterospecifics, such as vertical bars (Morris *et al*., 2005). This mixed evidenced on the role of female preferences in species where males have lost sexually selected traits leaves many unanswered questions about the drivers of this loss. Future work tackling whether changes in ornamentation in *X. pygmaeus* and *X. continens* are attributable to ecological shifts in these species (e.g. to inhabiting faster flowing rivers) or in part attributable to shifts in female preferences (e.g. weaker relative preferences) will shed light on the drivers of the repeated loss of male ornamentation.

More broadly, our results have implications for understanding convergent evolution at the phenotypic level. We find that suites of sexually selected ornaments—including the sword, the sword edge, and vertical bars—are coincidently lost with the evolution of small male body size. Available genetic mapping data for the sword and sword edge indicates that each of these traits is likely controlled by multiple loci (Powell *et al*., 2020). Similar repeated shifts in suites of traits have been previously reported in the context of adaptation to particular ecological conditions (Rennison *et al*., 2019) or to certain pollinators (Wessinger *et al*., 2019). Relative to convergent evolution of quantitative ecological traits, much less is known in practice about the drivers of convergent evolution of suites of sexually selected traits. Rigorously testing hypotheses about the drivers of convergent evolution of ornamentation require comparative studies of both potential ecological drivers and mate preferences across multiple species. Our results highlight the need for such studies in order to understand the recurrent gains or losses of suites of sexually selected traits in *Xiphophorus* and beyond.

## Acknowledgements

We thank Stepfanie Aguillon, Rongfeng Cui, Julia Palacios, Julie Zhang, Julia Palacios, and members of the Schumer lab for helpful discussion and/or feedback on earlier versions of this work. We thank Stanford University and the Stanford Research Computing Center for providing computational support for this project. This work was supported by a Hanna H. Gray, Pew and Searle fellowship to MS, the Ohio University Rush Elliot Professorship to MRM, and National Institute of Health R24-OD031467 to YL.

## Data and code availability

All code generated for or used in this project is available at https://github.com/Schumerlab/phylogeny_update and https://github.com/Schumerlab/Lab_shared_scripts. All raw data will be deposited on the NCBI SRA. All phenotypic data will be deposited on Dryad.

