## Supplementary Information for "Recurrent evolution of small body size and loss of the sword ornament in Northern Swordtail fish"

**Supplementary Information 1. Phenotypically misidentified *X. nigrensis* female and analysis of lab-maintained *X. nigrensis* sample**

The only northern swordtail species for which we were unable to acquire a wild-caught sample for sequencing was *X. nigrensis*. We initially hoped to sequence a recently wild-caught sample for this species using preserved DNA originally derived from the colony at UT Austin. No voucher photograph was available for this individual. Given the difficulty of obtaining *X. nigrensis* samples, we proceeded with sequencing the sample anyway and analyzed it as described in the main text for other northern swordtail samples.

We found that this sample clustered with *X. pygmaeus* (Fig. S2), in contrast to results from previous studies (Cui et al. 2013; Jones et al. 2013). Further investigation of this sample suggested that was likely an early generation hybrid between *X. pygmaeus* and *X. nigrensis.* Analyzing $D_{xy}$ between this sample and those of related species in windows, we saw that large regions of the genome had low $D_{xy}$ to *X. pygmaeus* samples whereas others had high $D_{xy}$ to *X. pygmaeus* and low $D_{xy}$ to *X. multilineatus* (*X. nigrensis'*s sister species based on previous analyses; Fig. S3). We thus suspect that this sample is an early generation hybrid between *X. pygmaeus* and *X. nigrensis* (or *X. multilineatus*). Consistent with this, $f_{branch}$ analysis highlighted substantial gene flow between this sample and *X. multilineatus* in an analysis where the genome-wide tree for this sample, *X. multilineatus*, *X. pygmaeus*, and outgroup northern swordtail species was provided (Fig. S1).

We sequenced another an individual from another lab colony descended from originally wild-caught *X. nigrensis* samples with voucher photos from at UT Austin. We performed whole genome sequencing of one male as described in the main text and repeated the analyses described above. This individual was inferred to be the sister group of *X. multilineatus*, as reported by previous phylogenies (Cui et al. 2013; Jones et al. 2013). The population this individual was derived from has been maintained in lab for ~15 generations but heterozygosity ($\pi$=0.13% per basepair) was similar to levels observed in high diversity species such as *X. birchmanni*, *X. variatus*, and *X. cortezi*. $f_{branch}$ analyses including this sample did not show evidence of substantial gene flow with other Northern swordtail samples, suggesting that it did not have a history of hybridization in captivity (Fig. 4).

**Supplementary Information 2. Branch length scaling for phylogenetically independent contrasts**

We faced two challenges in setting branch lengths for phylogenetically independent contrasts. The first challenge was appropriately scaling branch lengths from previous phylogenetic datasets for species not included in this study. In particular, although we had access through publicly available data or data collected for this project to whole-genome datasets for northern swordtails and platyfish, we lacked whole-genome data from most southern swordtails. We used the phylogeny developed by Cui *et al.* (2013) for species not included in our dataset. However, since that phylogeny was inferred from RNAseq data, the branch lengths will systematically differ from those we infer based on whole genome data. To address this, we performed analyses based on pairs of species for which we collected whole-genome resequencing data and were included in the Cui et al. study, and calculated the ratios of the branch lengths. We found that the ratios ranged from 10-25 (Table S6). We thus performed analyses using the median value to rescale branch lengths from Cui et al. for species we had not sampled in the present study. This allowed us to include data from the whole genus in PIC analysis. To evaluate the robustness of our results to the exact branch length rescaling factor used, we also rescaled branches using the minimum and maximum inferred ratios (Table S6), and re-ran PIC analysis for comparisons with body size that were significant in our main analysis (Tables S3-S4). We found that our results for this analysis were qualitatively unchanged.

The second challenge was determining how to include within-species variation. For *X. multilineatus* and *X. nigrensis*, where multiple male morphs exist within species, we needed to determine the appropriate branch lengths for these within-species comparisons. To do so, we wanted to measure the branch lengths separating individuals of the same species for both *X. multilineatus* and *X. nigrensis*. Since an individual's two haplotypes are samples of two parental individuals from the same population, we initially calculated $\pi$ per basepair for our *X. multilineatus* and *X. nigrensis* samples and used this measure as the branch length separating the large and small morphs within each species. However, in practice we found that these short branches resulted in strong correlations between the standardized independent contrasts and their standard deviations (following Garland 1992). Past work has indicated that such correlations have the potential to inflate the false positive rate in phylogenetically independent contrasts analysis (or PIC, see next section). We reasoned that given the phylogenetic distribution of traits of interest in the pygmy swordtail clade (i.e. ornamentation and body size), polymorphism for these traits likely existed in the common ancestor of *X. multilineatus* and *X. nigrensis*. We thus set the branch lengths for the small and large morphs equal to the branch length that separated *X. multilineatus* and *X. nigrensis*. We found that by using these branch lengths, correlations between the standardized independent contrasts and their standard deviations (Garland 1992) for body size and other traits were absent or much reduced.

Since the approach we implemented to account for polymorphism in *X. multilineatus* and *X. nigrensis* is not commonly used, we repeated the analysis using two approaches that did not require us to make assumptions about branch lengths within species. For all of these approaches, we included one tip for each *X. multilineatus* and *X. nigrensis* used the inferred branch length from the whole-genome RAxML phylogeny. In the first analysis, we excluded the small male morph from *X. multilineatus* and *X. nigrensis* and repeated PIC for the traits of interest identified in our main analysis. In the second analysis, we excluded the large male morph from both species. We found that despite reduced power, the results for sword presence versus absence, vertical bar number, PC1 of sexual dimorphism, dorsal index, and gonopodial ray 3a angle were significantly associated with body size in all analyses (Table S5).

**Supplementary Information 3. Tests for potential artifacts arising in PIC analyses**

Phylogenetically independent contrasts (PIC) analyses address the issue that species share ancestry and, as a result, their trait values are not independent. Thus, classic implementations of PIC transform the trait values of interest to account for expectations under this shared evolutionary history assuming an evolutionary model of Brownian Motion (Garland *et al.*, 1992; Díaz-Uriarte & Garland, 1996). In particular, phylogenetically independent contrasts methods compute weighted differences in a trait of interest between pairs of nodes based on the topology of the phylogeny. This results in *n-1* contrasts for a given *n* species included in the phylogeny. Barring errors in phylogenetic reconstruction, these contrasts are statistically independent and can be used downstream in standard analyses for correlations between traits of interest (e.g. linear regression analysis). However, statistical research and simulation studies have demonstrated that variation in branch lengths in the focal phylogeny can cause issues that increase the false positive rate in downstream analyses (Garland *et al.*, 1992; Díaz-Uriarte & Garland, 1996). In particular, contrasts inferred from longer branches (typically corresponding to longer periods of time) are likely to produce more extreme values and to have greater weight in analyses of the correlation between two variables. If unaddressed, this can produce an inflated false positive rate in these tests (Garland *et al.*, 1992). A solution to this issue is to standardize the independent contrasts so that they receive equal weighting in downstream analyses. Garland *et al.*, 1992 proposes plotting the absolute value of each independent contrast versus its standard deviation and evaluating whether there is a significant trend in the data. The presence of a significant trend indicates that the contrast of interest has not been effectively standardized, potentially leading to problems with an inflated false positive rate in analyses involving that contrast.

We performed these analyses for each continuous trait (i.e. contrast) of interest (Table S3). We found that several had significant trends between the absolute value of the independent contrast and the standard deviation (Fig. S10). For these traits, we attempted transformations of branch lengths to remove the significant trend (Fig. S10; Garland *et al.*, 1992). Description of any transformation applied to analyses of individual traits are provided in Table S3.

We also evaluated the expected performance of PIC applied to our dataset using simulation-based approaches. First, we evaluated whether the expected false positive rate matched the observed false positive rate using null simulations, where no correlated evolution with body size was modeled (using body size measurements obtained from the real data). We performed simulations for both continuous and binary traits. For continuous traits, we used the fastBM function (Revell, 2012) to simulate evolution along the swordtail phylogeny of traits with a mean trait value matching 1) the observed sword index, 2) the observed sword edge width, and 3) PC1 of sexual dimorphism. We set the bounds in each simulation to the observed range of trait values in our dataset, and heuristically set fastBM's sig2 parameter to produce simulations that approximated the observed trait distributions across the phylogeny in the real data. An example of this is shown in Fig. S11. For each scenario, we performed 1,000 simulations and calculated the p-value for co-evolution between body size and the simulated trait using the pic command, as we had for the real data. P-value distributions from these simulations are shown in Fig. S12. Based on these results, we see evidence of a slightly inflated false positive rate for some traits, but p-value distributions are relatively uniform (Fig. S12).

We next used a similar approach to simulate binary traits along the swordtail phylogeny using the function rbinTrait (Tung Ho & Ané, 2014). Here, we focused on simulating a trait with a similar distribution of presence and absence to the sword ornament when coded as a binary phenotype (Fig. S13). For each simulation, we tested the correlation with body size using the phylogenetic logistic regression approach of Ives and Garland (2010), as we had for the real data. We found that the p-value distribution from this analysis is expected to be conservative (Fig. S12).

We next explored our expected power using simulations of coevolution with the R command rTraitMult (Paradis & Schliep, 2019). Since the true parameters for coevolution between traits are unknown, we simply explored general scenarios to determine if we are likely to be underpowered given the number of species and timescales studied. rTraitMult allows users to set the root value and a covariance parameter (gamma) to model two traits and simulate their evolution along a phylogeny. We used a model of Brownian Motion for the evolution of continuous traits following Paradis (2014). For our initial simulations modeling coevolution between body size and sword length, we set the root values to the median male length and the median sword length (phylogenetic analyses suggest that the sword was present at the base of the genus; Cui *et al.*, 2013). We performed each simulation 1,000 times, varying gamma between 0.01 and 0.13. This range of gamma values generated phylogenetically corrected trait correlations ($R^2$) ranging from 0.08-0.82. For each gamma value, we determined the proportion of simulations where the simulated traits were significantly correlated at $p<0.05$ after PIC correction, using the same approach applied to the real data. We found that when gamma exceeded 0.06, we had good power to detect coevolution of these simulated traits. These results are summarized in Fig. S14.

**Supplementary Information 4. Demographic analyses and analyses of gene flow in Northern swordtails**

*PSMC results*

We used the pairwise sequentially Markovian coalescent or PSMC approach of Li and Durbin (2011) to infer changes in effective population size over time in wild caught samples of *Xiphophorus* species for which whole-genome sequence data had not been previously collected. With the caveat that comparing results across species makes strong assumptions that generation times and per-generation mutation rates have not evolved, results of these analysis points to unique demographic histories for each lineage (Fig. 4, Fig. S8).

*X. continens* and *X. montezumae* have the lowest genetic diversity ($\pi$) of the species analyzed (0.03%). While *X. montezumae* appears to have undergone a sustained population bottleneck starting around 10,000 generations before the present (assuming a generation time of 2/year), *X. continens* has maintained an estimated effective population size of ~20,000 individuals over a similar time period.

The pygmy swordtails *X. pygmaeus* and *X. multilineatus* have similar patterns of change in historical effective population size in the more distant past. We infer that *X. pygmaeus* experienced a reduction in population size around 10,000 generations before the present, followed by substantial population growth. Bootstrapping results indicate that we lose confidence in estimates for *X. pygmaeus* around this period of inferred population growth (Fig. S8). Results for *X. multilineatus* point to a more dramatic reduction in population size around the same period. However, this also falls within a time period where we lose confidence in our estimates for *X. multilineatus*.

An important note in comparing results across species is that our analysis explicitly assumes the same mutation rate and generation time for all species. In particular, the assumption of the same generation time is likely violated. Generation time varies substantial across *Xiphophorus*, and our own experiences in lab have suggested shorter generation times for *X. pygmaeus* than other northern swordtails. Little is known about the generation times of *X. continens* and *X. montezumae*. Thus, we avoid explicitly comparing trends across species in light of this uncertainty.

Besides basic knowledge about the population history of these species, a major goal of our PSMC analysis was determining whether species that have lost male ornamentation have undergone bottlenecks. Across species that have lost male ornamentation, we do not observe a consistent pattern of smaller historical effective population sizes, although we note that we lose resolution for all species in the last ~2,000-10,000 generations. Although we cannot rule out the possibility that genetic drift played a role in the loss of some ornaments in the recent past, we note that we observe directional losses of all ornamentation traits in *X. continens* and *X. pygmaeus*, as opposed to a combination of fixation and loss, as would be expected under a drift model.

*Analysis of gene flow using Dsuite*

Past work using coalescent and site-based methods revealed rampant gene flow between *Xiphophorus* species, including in the Northern swordtail clade (Cui *et al.*, 2013). With newly available whole-genome sequences for *X. continens*, we wanted to revisit these patterns. Due to the computational intensity of coalescent based methods (Liu *et al.*, 2009), especially for whole-genome sequences, we focus on site-based analyses, noting cases where the accuracy of our inferences might be expected to be reduced.

As described in the main text (see Methods), we used the package Dsuite to infer possible admixture between species in the Northern swordtail clade. We found evidence of substantial gene flow between extant species as well as between lineages ancestral to modern day species (Fig. 4, Fig. S9). In our analysis of all species in the Northern swordtail clade, we see signals of gene flow between *X. cortezi* and nearly every other species in the clade. *X. cortezi* is broadly distributed, with range overlap with *X. pygmaeus, X. multilineatus*, and *X. birchmanni* (Fig. 4), and has active hybrid zones with *X. birchmanni* (Powell *et al.*, 2021; Langdon *et al.*, 2022). Its range also abuts that of *X. montezumae* and *X. nezahualcoyotl* and indeed *X. nezahualcoyotl* is known to be an ancient hybrid between *X. montezumae* and *X. cortezi* (Schumer *et al.*, 2016). We note that simulations in the original Dsuite paper suggest that sometimes $f_{branch}$ signals can be erroneously inflated with close relatives of the hybridizing species, so it is not clear from the current analyses that *X. cortezi* has hybridized with all of the species discussed above. However, many of these signals of ancient and contemporary gene flow with *X. cortezi* were detected in previous analyses using RNAseq data from Northern swordtails (Cui *et al.*, 2013). For simplicity, we present analyses including *X. cortezi* in Fig. S9 but exclude these results from the main text.

Focusing on the clades within Northern swordtails where ornamentation has been fully or partially lost, the pygmy swordtails and *X. continens*, we can see a number of interesting patterns, but with no obvious connections to the loss of ornamentation (Fig. 4). We see evidence of substantial gene flow with *X. nezahualcoyotl* (~10% of the genome) but weak evidence for genetic exchange with other species. As *X. nezahualcoyotl* is an ornamented, large-bodied species (Fig. 2), this genetic exchange is unlikely to be linked to the loss of ornamentation.

*PhyloNetHMM analysis*

Dsuite analysis suggests that there has been extensive gene flow between *X. continens* and *X. nezahualcoyotl*. We were interested in further investigating the likely direction of that gene flow. Given the sister relationship between *X. continens* and *X. montezumae*, we reasoned that if haplotypes originated in *X. nezahualcoyotl* and introgressed into *X. continens*, these regions would be expected to have lower sequence divergence between *X. nezahualcoyotl* and *X. continens* but high sequence divergence between *X. continens* and *X. montezumae* compared to other regions of the genome. By contrast, if they originated from *X. continens* and spread into *X. nezahualcoyotl*, divergence between *X. continens* and *X. montezumae* in these regions is predicted to be unremarkable, and both *X. continens* and *X. montezumae* should have lower divergence from *X. nezahualcoyotl* within these regions.

We used the phylogenetic local ancestry inference tool PhyloNetHMM (Liu *et al.*, 2014) to identify regions of the genome that show signals consistent with admixture between *X. continens* and *X. nezahualcoyotl*. We analyzed whole genome alignments that included *X. continens*, *X. montezumae*, *X. nezahualcoyotl*, and used *X. variatus* as the outgroup sequence. We used a posterior probability cutoff of 0.9 for identifying regions of the genome as matching the "species tree" (i.e, *X. continens* and *X. montezumae* grouped as sister taxa) or the "gene flow tree" (i.e. *X. continens* and *X. nezahualcoyotl* grouped as most closely related). PhyloNetHMM attempts to model ILS by allowing gene trees to be discordant, and earlier simulation studies suggest that it is relatively accurate at distinguishing ILS from gene flow (Schumer *et al.*, 2016).

However, some of the tracts identified in our analysis may group *X. continens* and *X. nezahualcoyotl* due to sequence similarity from ILS.

67% of the genome fell into regions that had a posterior probability of >0.9 of either the species tree or the gene flow tree. We analyzed pairwise sequence divergence ($D_{xy}$) between *X. continens-X. montezumae*, *X. continens-X. nezahualcoyotl*, and *X. montezumae-X. nezahualcoyotl* separately for the species tree and gene flow tree tracts. The results of this analysis are shown in Fig. S15. These patterns are inconsistent with our expectations under a model of unidirectional gene flow from *X. nezahualcoyotl* into *X. continens*, and we interpret them as preliminary evidence that genetic exchange occurred primarily from *X. continens* into *X. nezahualcoyotl.* However, we interpret these results with caution since the proportion of called regions inferred to have been exchanged between *X. nezahualcoyotl* and *X. continens* based on PhyloNetHMM analysis far exceeds the estimate from Dsuite analysis (34% of called regions vs Fig. 4).

Both analyses with Dsuite and PhyloNetHMM support substantial genetic exchange between *X. nezahualcoyotl* and *X. continens*. This is particularly notable in understanding the evolutionary history of *X. nezahualcoyotl* because of previous findings. Past work has found substantial genetic exchange between *X. cortezi* and *X. nezahualcoyotl* (Schumer *et al.*, 2016), highlighting a complex history of hybridization in the *X. nezahualcoyotl* lineage.

**Supplementary Information 5. Additional phenotypic analyses of body size**

*Collecting data from the literature on body size and re-analyzing phylogenetically corrected correlations with body size*

While measuring a single individual of each species provides a relatively unbiased estimate of body size, it also introduces substantial noise from within species variation that may reduce our power to detect relationships between body size and other traits of interest. Thus, we collected data from the literature on average male length for species for which this data was available. When searching for standard length information across species, the keywords 'adult standard length,' 'body size,' and '*Xiphophorus* size measurement' were used along with each species name. The articles were then sorted based on applicability. Criteria for inclusion of data from the literature were data from reproductively mature males and females either measured from specimens collected directly from natural populations or from individuals collected from natural populations and maintained in the lab. We also required that the studies reported either raw data or average length of individuals, the sample size used, and excluded studies where measurements were taken from fewer than three adult individuals. Data for each species is reported in Table S7. For the following species, we were unable to collect this data, data included fewer than three individuals, or the sample size was not reported in the literature: for both males and females of *X. alvarezi, X. signum*, *X. mayae*, *X. kallmani* and females of *X. andersi* and *X. milleri.* For those cases, we used our initial measurements.

Since this data was collected over the past 60 years in a variety of scenarios, there is significant variation in the methods and sampling strategy reflected in this literature-based dataset. We thus approach the interpretation of these analyses with caution, but look to confirm the general patterns we observed in PIC analyses with our primary dataset. Reanalyzing this dataset as described in the main text, we found that traits that were significantly correlated with body size in our initial analysis, with the exception of dorsal index, remained correlated or showed a trend in the same direction as our original analysis (Table S8).

*Quantifying population-level body size for species reported to have male body size polymorphism*

We measured standard length for two species (*X. variatus* and *X. nezahualcoyotl*) reported to be polymorphic for male body size in the literature but for which body size polymorphism has not been well-documented. We also collected data for *X. multilineatus* and *X. nigrensis*, species where multiple morphs are well-documented in the literature, and for a Northern swordtail species without body size polymorphism (*X. cortezi*). Standard length for each individual included in the dataset was measured from photos in ImageJ. Distributions were visualized with the R package ggjoy. To statistically test for evidence that the distribution of data did not fit a unimodal distribution, we employed Hartigan's test for multimodality with the R package diptest (dip.test() with default parameters). The results for each species analyzed are shown in Figure 2.

**Supplementary Information 6. Converting sword index and related measurements into a binary variable for analysis of binary covariates**

Several approaches for PIC allow researchers to compare coevolution of binary traits and continuous traits. In some cases where continuous traits have a bimodal distribution, it is useful to convert these traits into binary variables (Ives & Garland, 2010). The sword index (the length of the sword ornament divided by an individual's body length) is one such trait (Fig. S6).

Based on the distribution of sword index, there are several points at which a line could be drawn to convert the sword index from a continuous into a binary trait. We tested three points that seemed reasonable based on the distribution. In the main text we present the analysis where individuals with >0.01 sword length are treated as sworded. Here, we explore more lenient cutoffs where individuals with short swords (sword indices of 0.014 and 0.05 respectively, are treated as unsworded; purple and blue dashed lines in Fig. S6).

With these thresholds, our results are qualitatively similar to those presented in the main text albeit less significant. With a threshold of 0.014, we find similar results (Log Likelihood = -11, p=0.007). With a threshold of 0.05, the relationship between sword presence and body size is marginally significant (Log Likelihood = -14, p=0.044).

Other traits that have a bimodal distribution in our data include the black pigmented edge of the sword ornament. Most species with a sword have a pigmented lower edge, but only some species have a pigmented upper sword edge. The length of the pigmented stripe on the upper and lower edge are very strongly correlated with sword length ($R_{sword\text{-}lower}$=0.86; $R_{sword\text{-}upper}$=0.71) so we did not analyze these characteristics as separate traits. However, the width of both the upper and lower edges varies dramatically between species (e.g. compare *X. montezumae* to *X. multilineatus* in Figure 2 of the main text). We thus collected width measurements for all species. Visualizing these width measurements (Fig. S6). Because the data had a somewhat bimodal distribution, we also analyzed these variables as binary variables, based on whether the species had an upper or lower sword edge width greater than zero (Fig. S6). When analyzed as a binary trait, neither upper or lower edge was significantly associated with body size (Table S4).

**Supplementary Tables**

**Table S1.** Details on where information on the lab strains or collection sites of Northern Swordtail samples used in previous phylogenies of *Xiphophorus* can be found in the original publications.

| **Publication** | **Localities** | **Source data in the original paper** |
|---|---|---|
| Meyer *et al.,* 1994 | Unavailable | Unavailable |
| Morris *et al.,* 2001 | Multiple | Appendix 1 |
| Kang *et al.,* 2013 | Multiple | Additional file 6 |
| Cui *et al.,* 2013 | Multiple | Table S1 |

*Table S2. The phenotypic dataset collected for PIC as part of this study is provided as an attached excel document.*

**Table S3.** Correlations between body size and all continuous traits analyzed using phylogenetically independent contrasts. Analyses were performed using the *pic* command in the *ape* R package. Where relevant, data transformations used or unavailable data is noted in the final column. PIC – phylogenetically independent contrasts. P-values and correlation coefficients are based on regression through the origin.

| **Trait 1** | **Trait 2** | **Diagnostic analysis p-value for Trait 2 (post-transformation p-value)** | **PIC correlation coefficient (R)** | **PIC P-value (Bonferroni)** | **Notes** |
|---|---|---|---|---|---|
| Body size | Sword index | 0.013 (0.07) | 0.42 | 0.028 (0.17) | Failed initial diagnostic test, branch lengths squared for analysis |
| Body size | Lower sword edge width | 0.041 (0.183) | 0.39 | 0.045 (0.27) | Failed initial diagnostic test, branch lengths squared for analysis |
| Body size | PC1 of sexual dimorphism analysis | 0.6 | 0.72 | 0.000021 (0.00013) | |
| Body size | PC2 of sexual dimorphism analysis | 0.11 | 0.37 | 0.056 (0.34) | |
| Body size | Vertical bar number | 0.56 | 0.70 | $4.3 \times 10^{-5}$ (0.0002) | |
| Body size | Dorsal index | 0.018 (0.10) | 0.65 | 0.00024 (0.001) | Failed initial diagnostic test, branch lengths squared for analysis |

**Table S4.** Correlations between body size and all binary traits analyzed using a phylogenetic logistic regression approach. Analyses were performed using the *phylolm* package with the logistic_IG10 method. Where relevant, unavailable data is noted in the final column.

| **Trait 1** | **Trait 2** | **Log Likelihood** | **Penalty Log Likelihood** | **PIC P-value (Bonferroni)** | **Notes** |
|---|---|---|---|---|---|
| Body size | Sword presence or absence | -11.0 | -8.2 | 0.0067 (0.027) | |
| Body size | Lower sword edge presence or absence | -13.1 | -10.4 | 0.082 (0.33) | |
| Body size | Upper sword edge presence or absence | -13.4 | -10.5 | 0.052 (0.21) | |
| Body size | Gonopodial spine angle ray 3 | -13.2 | -9.9 | 0.032 (0.13) | *P. jonseii* excluded due to lack of data |

**Table S5.** Results of analyses exploring robustness of PIC results to including only one phenotypic group (large or small morph males) for *X. nigrensis* and *X. multilineatus*.

| **Trait 1** | **Trait 2** | **Analysis version** | **PIC correlation coefficient (R) or Log Likelihood** | **PIC P-value** |
|---|---|---|---|---|
| Body size | Sword index | Exclude large | 0.27 | 0.24 |
| | | Exclude small | 0.35 | 0.088 |
| Body size | Lower sword edge width | Exclude large | 0.16 | 0.45 |
| | | Exclude small | 0.26 | 0.2 |
| Body size | PC1 of sexual dimorphism | Exclude large | 0.78 | $4.4 \times 10^{-5}$ |
| | | Exclude small | 0.71 | $7.9 \times 10^{-4}$ |
| Body size | Vertical bar number | Exclude large | 0.75 | $1.6 \times 10^{-5}$ |
| | | Exclude small | 0.69 | 0.00016 |
| Body size | Dorsal index | Exclude large | 0.64 | 0.0006 |
| | | Exclude small | 0.63 | 0.0008 |
| Body size | Sword presence or absence | Exclude large | -8.0 | 0.011 |
| | | Exclude small | -7.2 | 0.009 |
| Body size | Gonopodial spine angle ray 3 | Exclude large | -9.7 | 0.022 |
| | | Exclude small | -6.5 | 0.010 |

**Table S6.** Ratios of branch lengths inferred by Cui et al. 2013 and the present study for one or more pairs of species for each major clade. Cui et al. 2013 used coding sequences only so branch lengths are expected to be systematically lower than those inferred from whole genome sequences.

| **Clade 1** | **Clade 2** | **Branch length – Cui et al. 2013** | **Branch length – present study** | **Ratio** |
|---|---|---|---|---|
| *X. clemenciae* | *X. hellerii* | 0.0127 | 0.323 | 25.4 |
| *X. birchmanni* | *X. malinche* | 0.0041 | 0.074 | 18.1 |
| *X. meyeri* | *X. gordoni* | 0.0013 | 0.0136 | 10.4 |
| *X. variatus* | *X. evelynae* | 0.0059 | 0.126 | 21.4 |
| *X. pygmaeus* | *X. multilineatus* | 0.0034 | 0.051 | 14.9 |

**Table S7.** Standard length measurements collected from the literature. Standard length measured from the anterior most point of the fish to the posterior most point of the hypural plate was collected from the literature for both males and females of each species. Data was gathered for average standard length and number of individuals measured. Sources that reported data for fewer than three individuals or that did not report the number of individuals measured (entries in red colored text) are noted here but were not used in our analysis. M – male, F – female.

| Species | Sex | Average Standard Length | Number Measured | Publication |
|---|---|---|---|---|
| *X. alvarezi* | M | 39.9 mm | 2 | Culumber & Tobler (2018) |
| *X. alvarezi* | F | 39.8 mm | 2 | Culumber & Tobler (2018) |
| *X. andersi* | M | 34 mm | 8 | Culumber & Tobler (2018) |
| *X. andersi* | F | 55 mm | Unavailable | Wischnath (1993) |
| *X. birchmanni* | M | 49.3 mm | 8 | Culumber & Tobler (2018) |
| *X. birchmanni* | F | 40 mm | 46 | Kindsvater (2012) Figure 1 |
| *X. clemenciae* | M | 44.4 mm | 14 | Kallman et al. (2004) Table 2 |
| *X. clemenciae* | F | 42.0 mm | 9 | Kallman et al. (2004) Table 1 |
| *X. cortezi* | M | 38.5 mm | 50 | Rosen (1960) |
| *X. cortezi* | F | 34.9 mm | 50 | Rosen (1960) |
| *X. couchianus* | M | 40 mm | 66 | Rosen (1960) |
| *X. couchianus* | F | 55 mm | 75 | Rosen (1960) |
| *X. continens* | M | 20.5 mm | 6 | This study |
| *X. continens* | M | 21-26 mm | 20 | Morris et al. (2005) |
| *X. continens* | F | 26-31 mm | 12 | Morris et al. (2005) |
| *X. evelynae* | M | 33 mm | 13 | Rosen (1960) |
| *X. evelynae* | F | 36 mm | 14 | Rosen (1960) |
| *X. gordoni* | M | 23.7 mm | 11 | Miller and Mickley (1963) |
| *X. gordoni* | F | 24.1 mm | 11 | Miller and Mickley (1963) |
| *X. hellerii* | M | 43.8 mm | 18 | Kallman et al. (2004) Table 2 |
| *X. hellerii* | F | 39.3 mm | 12 | Kallman et al. (2004) Table 1 |
| *X. kallmani* | M | - | Unavailable | |
| *X. kallmani* | F | - | Unavailable | |
| *X. maculatus* | M | 30.5 mm | 9 | Culumber & Tobler (2018) |
| *X. maculatus* | F | 34.5 mm | 8 | Culumber & Tobler (2018) |
| *X. malinche* | M | 45.7 mm | 12 | Rauchenberger et al. (1990) |
| *X. malinche* | F | 44.5 mm | 18 | Tudor & Morris (2008) |
| *X. mayae* | M | 53.6 mm | 1 | Meyer & Schartl (2002) |
| *X. mayae* | F | 61.8 mm | 1 | Meyer & Schartl (2002) |
| *X. meyeri* | M | 29 mm | 25 | Schartl & Schröder, (1988) |
| *X. meyeri* | F | 28.5 mm | 25 | Schartl & Schröder, (1988) |
| *X. milleri* | M | 24.1 mm | 8 | Culumber & Tobler (2018) |
| *X. milleri* | F | 28.5 mm | 1 | Rosen (1960) |

| | | | | |
|---|---|---|---|---|
| *X. montezumae* | M | 39.5 mm | 78 | Kallman (1983) |
| *X. montezumae* | F | 31.5 mm | 40 | Kallman (1983) |
| *X. montezumae* | M | 30 mm | >50 total M/F (some juveniles) | Rosen (1960) |
| *X. montezumae* | F | 30 mm | >50 total M/F (some juveniles) | Rosen (1960) |
| *X. monticolus* | M | 44.3 mm | 12 | Kallman et al. (2004) |
| *X. monticolus* | F | 49.2 mm | 9 | Kallman et al. (2004) |
| *X. multilineatus* | M | 23.7 mm sneaker<br>37.6 mm courter | 122 | Zimmerer & Kallman (1989) |
| *X. multilineatus* | F | 26 mm | 27 | Rios-Cardenas et al. (2013) |
| *X. nezahualcoyotl* | M | 29 mm | 43 | This study |
| *X. nezahualcoyotl* | F | 42-48 mm | 18 | Rauchenberger et al. (1990) |
| *X. nigrensis* | M | 24.1 mm sneaker<br>35.5 mm courter | 362 | Zimmerer & Kallman (1989) |
| *X. nigrensis* | F | 18.2 mm | 8 | Culumber & Tobler (2018) |
| *X. pygmaeus* | M | 22.2 mm | 8 | Culumber & Tobler (2018) |
| *X. pygmaeus* | F | 26.0 mm | 7 | Culumber & Tobler (2018) |
| *X. variatus* | M | 32.9 mm | 7 | Culumber & Tobler (2018) |
| *X. variatus* | F | 34.3 mm | 8 | Culumber & Tobler (2018) |
| *X. xiphidium* | M | 29.5 mm | 8 | Culumber & Tobler (2018) |
| *X. xiphidium* | F | 30.03 mm | 9 | Culumber & Tobler (2018) |
| *X. signum* | M | - | Unavailable | |
| *X. signum* | F | - | Unavailable | |

**Table S8.** Results of re-analysis of significant correlations using dataset composed of body size values reported in the literature.

| Trait 1 | Trait 2 | PIC correlation coefficient (R) or Log Likelihood | PIC P-value | Notes |
|---|---|---|---|---|
| Body size (literature) | Sword index | 0.39 | 0.045 | Branch lengths squared for analysis |
| Body size (literature) | Lower sword edge width | 0.35 | 0.071 | Branch lengths squared for analysis |
| Body size (literature) | PC1 of sexual dimorphism analysis | 0.33 | 0.09 | |
| Body size (literature) | Vertical bar number | 0.34 | 0.086 | |
| Body size (literature) | Dorsal index | -0.32 | 0.10 | |
| Body size (literature) | Sword presence or absence | -7.6 | 0.0062 | |
| Body size (literature) | Gonopodial spine angle ray 3 | -9.0 | 0.016 | *P. jonseii* excluded due to lack of data |

**Table S9.** PCA loadings for sexual dimorphism analysis presented in Figure 2A, for the first five PCs.

| **Trait** | PC1 | PC2 | PC3 | PC4 |
|---|---|---|---|---|
| standardized sword length | 0.04445 | -0.16219 | -0.02518 | 0.05324 |
| macromelanocyte pigment | 0.00031 | -0.02758 | -0.13816 | -0.02719 |
| micromelanocyte pigment | 0.04738 | -0.11925 | -0.90597 | -0.34700 |
| vertical bar number | 0.99499 | -0.02474 | 0.02663 | 0.06556 |
| caudal fin pigmentation | -0.05543 | -0.11588 | -0.23595 | 0.53863 |
| dorsal fin pigmentation | -0.03618 | 0.02320 | -0.24675 | 0.44523 |
| sword pigmentation | -0.01714 | -0.21948 | -0.09076 | 0.59748 |
| upper sword edge width | -0.03170 | -0.76746 | 0.18165 | -0.13576 |
| standardized upper sword edge length | -0.00027 | -0.00561 | -0.00231 | 0.00034 |
| lower sword edge width | -0.00688 | -0.55392 | 0.03146 | -0.08519 |
| standardized lower sword edge length | 0.00079 | -0.01026 | -0.00132 | 0.00630 |
| standardized dorsal fin length | -0.00011 | 0.00085 | 0.00048 | 0.00143 |
| standardized dorsal fin height | 0.00025 | 0.00058 | 0.00009 | 0.00048 |
| standardized body depth | -0.00002 | 0.00085 | 0.00103 | 0.00159 |
| standardized peduncle depth | -0.00012 | 0.00067 | 0.00027 | 0.00081 |
| standardized caudal fin length | 0.00000 | 0.00078 | 0.00039 | 0.00013 |
| standardized caudal fin height | 0.00607 | 0.00149 | -0.00028 | 0.02554 |

**Table S10.** PCA loadings for male phenotypic variation analysis for different species presented in Figure 2B, for the first five PCs.

| Trait | PC1 | PC2 | PC3 | PC4 | PC5 |
|---|---|---|---|---|---|
| Standard length | 0.2075 | 0.2918 | -0.1062 | -0.4860 | 0.1481 |
| Sword length | 0.4830 | -0.1306 | -0.1787 | 0.3635 | 0.4194 |
| Vertical bar number | 0.0487 | 0.2603 | -0.2208 | 0.1535 | 0.6331 |
| Upper sword edge presence | 0.0109 | -0.0228 | -0.0085 | -0.0228 | 0.0023 |
| Upper sword edge width | 0.0051 | -0.0124 | -0.0015 | -0.0073 | -0.0008 |
| Upper sword edge length | 0.3216 | -0.7638 | -0.1059 | -0.4470 | 0.0819 |
| Lower sword edge presence | 0.0109 | 0.0196 | 0.0039 | -0.0217 | -0.0894 |
| Lower sword edge width | 0.0090 | -0.0028 | 0.0211 | -0.0183 | -0.0094 |
| Lower sword edge length | 0.4554 | 0.1096 | 0.8213 | -0.0156 | 0.0300 |
| Dorsal fin length | 0.0761 | 0.1594 | 0.0722 | -0.2159 | 0.0778 |
| Dorsal fin height | 0.0435 | 0.0597 | 0.0240 | -0.0963 | -0.0254 |
| Body depth | 0.0614 | 0.1737 | 0.0909 | -0.1866 | 0.1901 |
| Peduncle edge | 0.0599 | 0.1701 | 0.0129 | -0.1345 | 0.1573 |
| Caudal fin length | 0.0625 | 0.1582 | -0.0372 | -0.2349 | -0.0564 |
| Caudal fin length | 0.0872 | 0.2936 | -0.2263 | -0.3494 | -0.2157 |
| Peduncle edge length | 0.0813 | 0.0310 | -0.2383 | -0.2066 | 0.0851 |
| Lower edge - peduncle length | 0.6139 | 0.1825 | -0.3134 | 0.2792 | -0.5110 |

### Supplementary figures

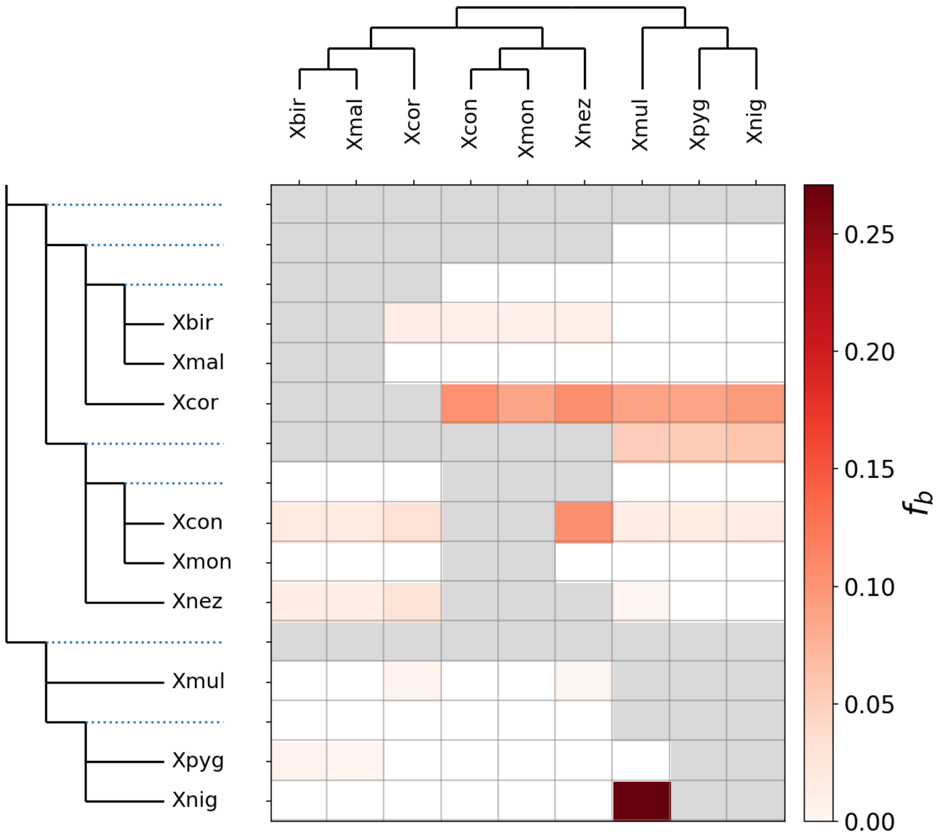


**Fig. S1.** Results of Dsuite analysis when the sample originally misidentified as *X. nigrensis* was included. Analyses of this sample showed evidence of recent hybridization with *X. pygmaeus* (Fig. S3). The $f_{branch}$ analysis shown here highlights the mixed ancestry in this individual since it is grouped with *X. pygmaeus* and is inferred to have substantial gene flow with *X. multilineatus*, the true sister species of *X. nigrensis*.

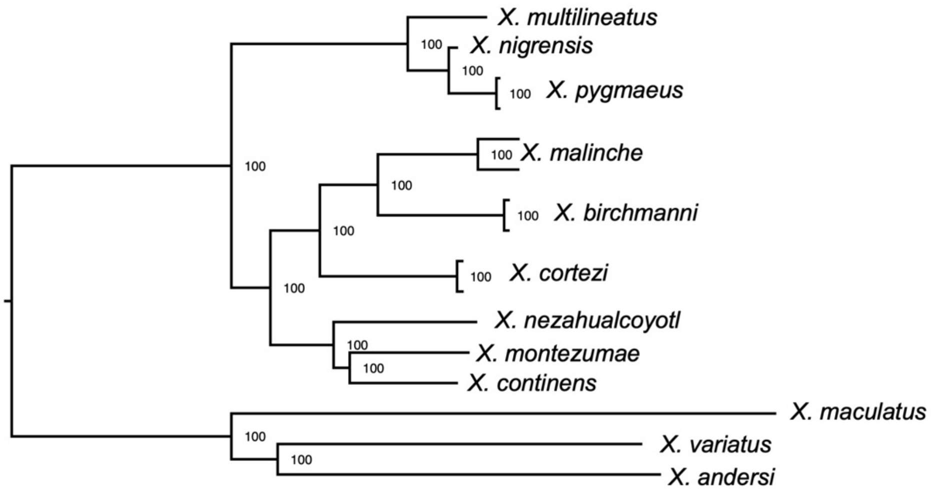


**Fig. S2.** RAxML phylogeny based on the misidentified *X. nigrensis* individual. Later analysis suggested that this individual was a hybrid between *X. nigrensis* and *X. pygmaeus*.

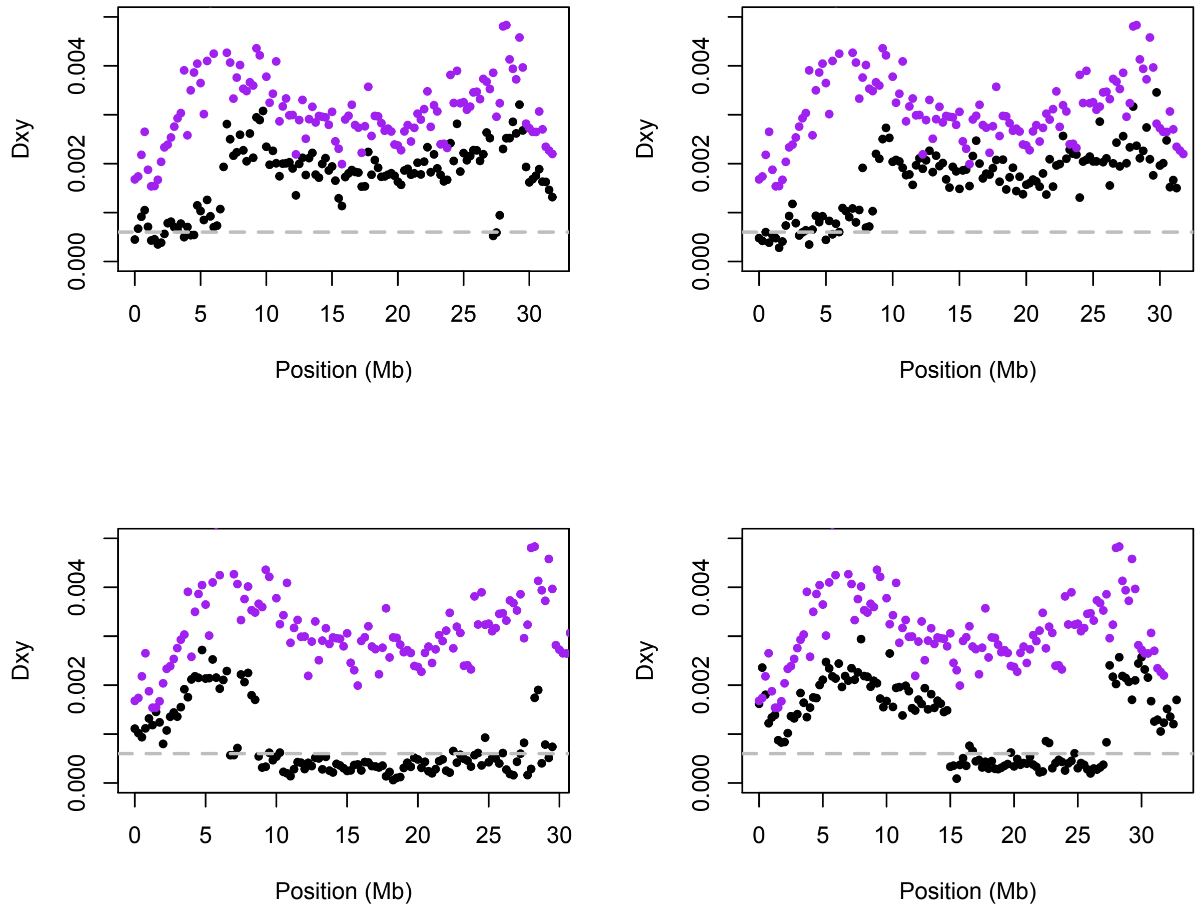


**Fig. S3.** Pairwise sequence divergence in 250 kb windows between *X. pygmaeus* and the putative hybrid sample (black) versus *X. pygmaeus* and the *X. nigrensis* lab strain used for the major analysis in the main text (purple). The gray dashed line indicates expected within-species polymorphism for *X. pygmaeus*. Four representative chromosomes are shown. The alternating regions of low and high divergence from *X. pygmaeus* in the putative hybrid sample are consistent with ancestry tracts from *X. pygmaeus* and another species (presumably *X. nigrensis*) in this sample. Thus, this individual was excluded from further analysis.

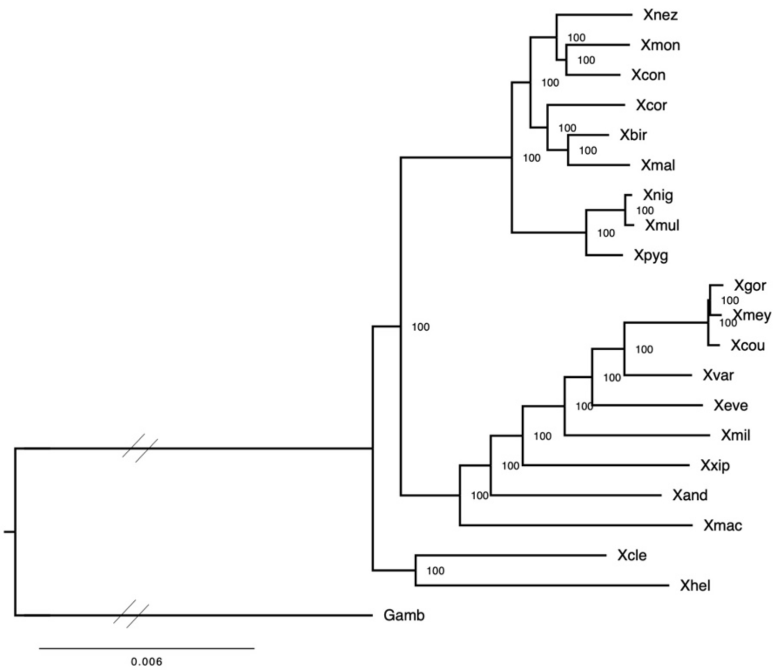


**Fig. S4.** Phylogenetic relationships inferred using RAxML from an alignment that included both variant and invariant sites for a subset of the genome. To generate a computationally tractable dataset for this analysis, we randomly subsampled 150 Mb from our whole genome alignment in 100 kb windows (approximately 20% of the genome), removed columns with missing data, and performed analysis with RAxML as described in the main text. Nodal support was estimated using 100 rapid bootstraps. Tree was rooted by the branch leading to *Gambusia affinis*. Angled lines on the phylogeny indicate branches shorted for visualization.

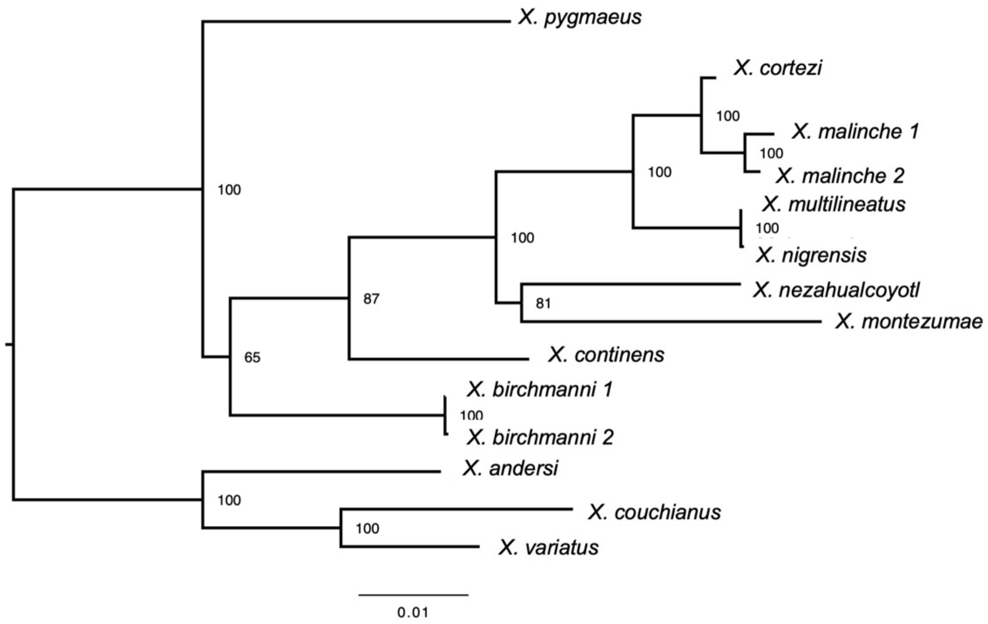


**Fig. S5.** Mitochondrial phylogeny for northern swordtail species produced by full mitochondrial sequence alignment using RAxML and the GTR + T model. Nodal support was generated with 100 rapid bootstraps using GTR + CAT. The topology was rooted by the branch to the platyfish lineages (*X. variatus*, *X. maculatus*, *X. andersi*).

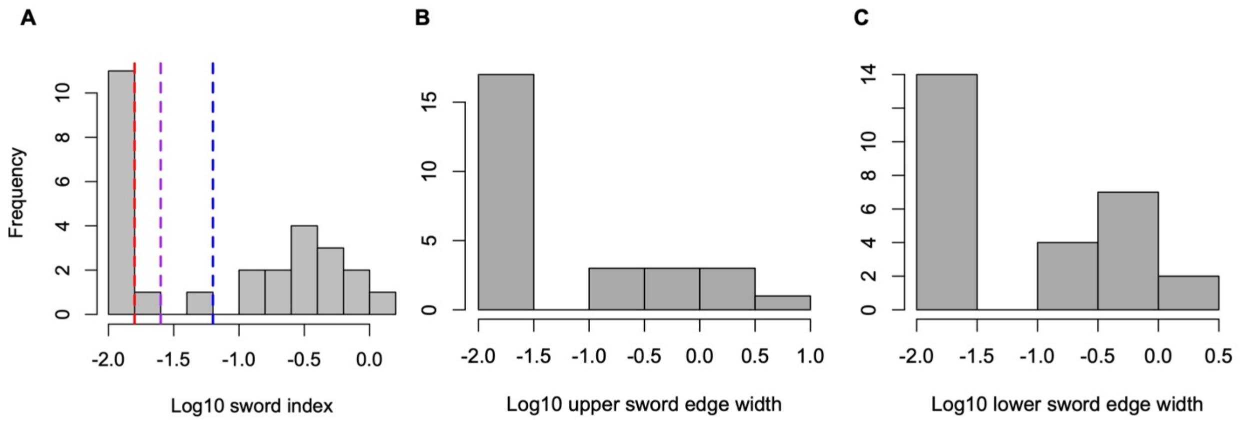


**Fig. S6. A)** Distribution of sword index, or male sword length divided by standard length, across *Xiphophorus* species. Colored lines show different thresholds used for calling a species sworded or unsworded in analyses presented in the main text and supplement. For the main text analyses, we treated all males with a sword index greater than zero as sworded (red line). **B**) Distribution of the upper sword edge width across *Xiphophorus* species. All males with an upper sword edge width greater than zero were treated as having the upper sword edge present. **C**) Distribution of the lower sword edge width across *Xiphophorus* species. All males with a lower sword edge width greater than zero were treated as having the lower sword edge present. Across all panels, the $Log_{10}$ of the trait of interest is plotted for visualization. For individuals with values of zero (i.e. a sword index of zero), 0.01 was used as their trait value for purposes visualization only.

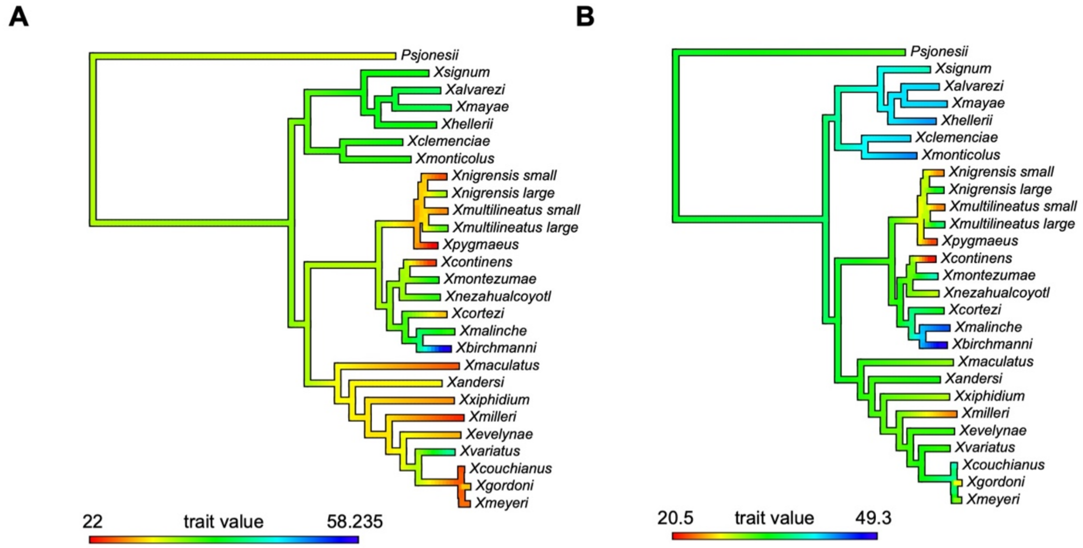


**Fig. S7.** Visualization of results for ancestral state reconstruction of body size, implemented in phytools. Ancestral states were estimated with the command fastAnc for male body size measurements collected for this paper (**A**) as well as average body size estimates collected from the literature (**B**). For both datasets, the ancestor of Northern swordtails is estimated to be 33-34 mm, however the 95% confidence intervals are large (**A** - 22-45 mm and **B** – 22-51 mm).

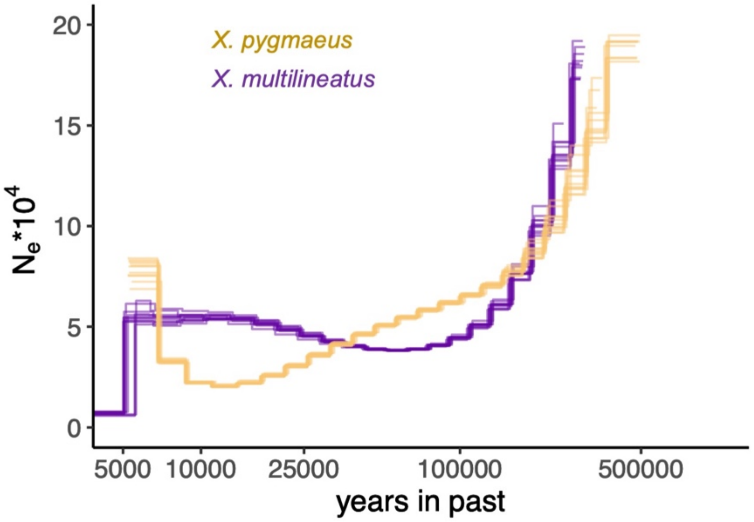


**Fig. S8.** Effective population size over time inferred from one *X. pygmaeus* and one *X. multilineatus* genome using PSMC. Analysis assumed a ratio of ρ/θ of 2, which matches the empirically inferred ratio for *X. birchmanni* (Schumer *et al.*, 2018), a mutation rate of 3.5 x $10^{-9}$ per basepair/per generation, and a generation time of 2 per year. Note that the generation time of *Xiphophorus* species in the lab differs.

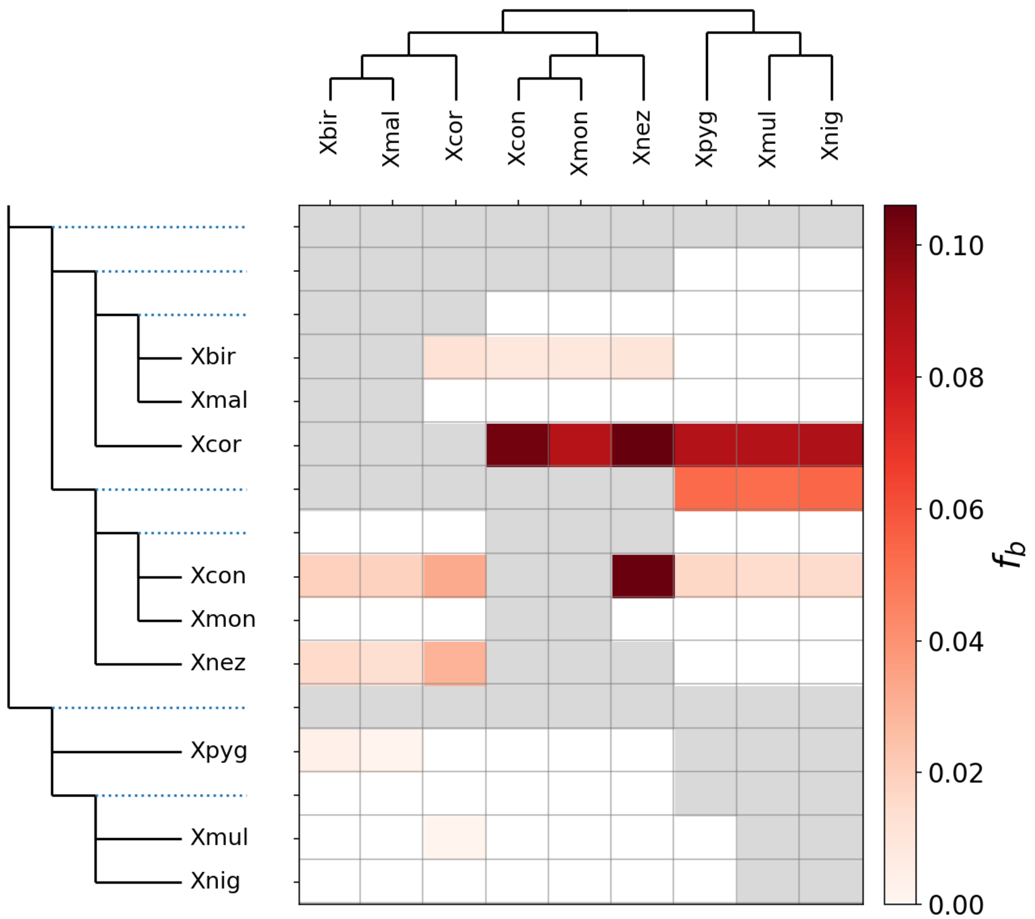


**Fig. S9.** Results of Dsuite analysis including *X. cortezi*, which has substantial evidence of gene flow with most species in the Northern swordtail clade. Some of these historical admixture events have been reported in previous studies of the group (Cui *et al.*, 2013). See Supporting Information 4 for discussion of these results.

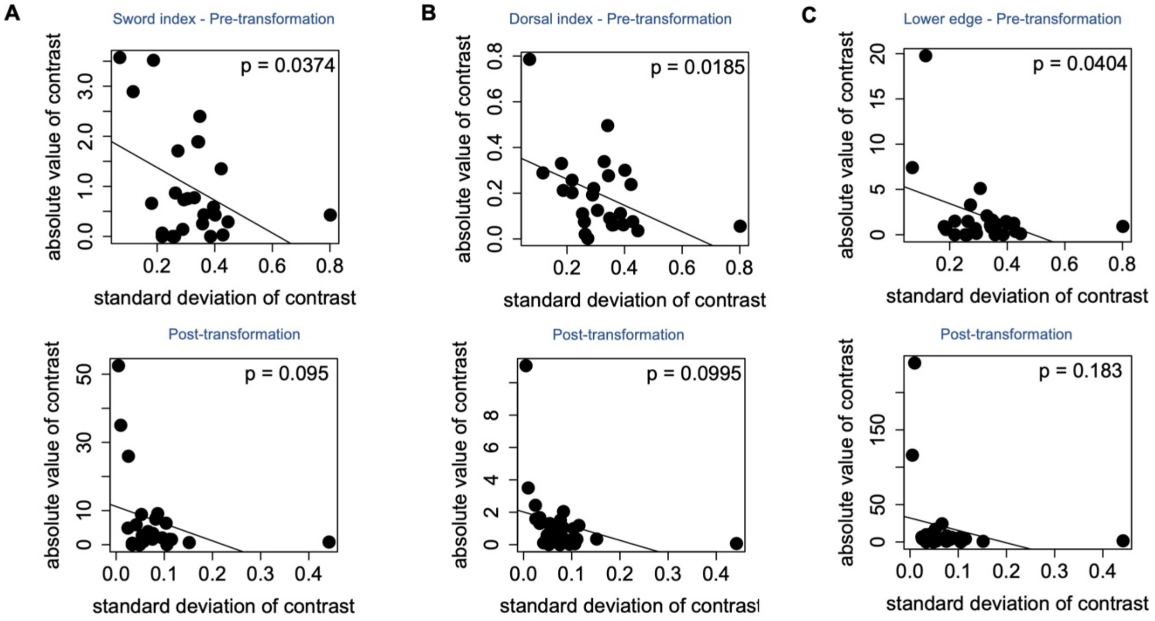


**Fig. S10.** Example diagnostic tests (Garland 1992) before and after branch length transformation for **A**) the sword index, **B)** the dorsal fin index and **C**) the width of the sword lower edge pigmentation.

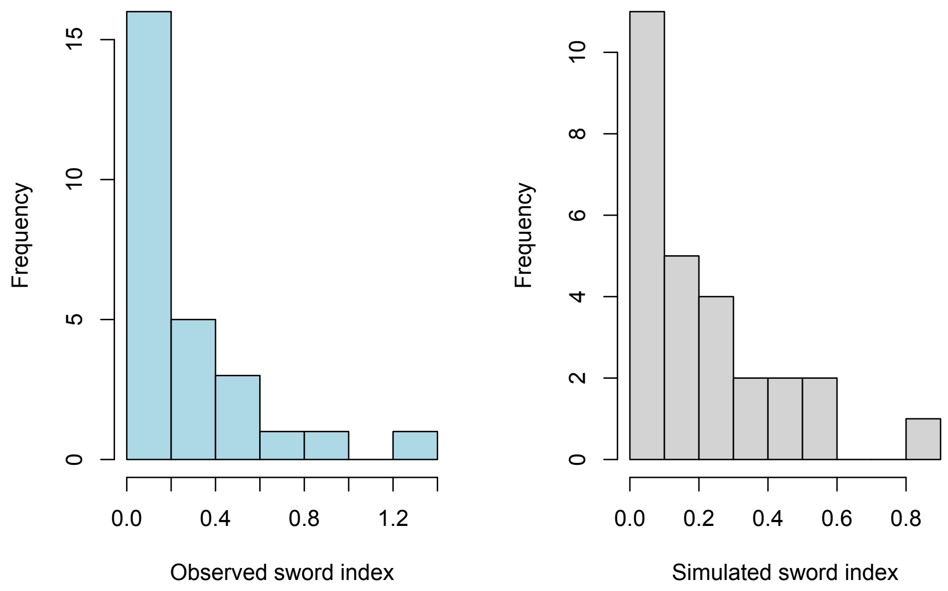


**Fig. S11.** Comparison of observed distribution of sword index in the *Xiphophorus* phylogeny to simulations of the sword index using the fastBM program in R. We used these simulations to evaluate the performance of the PIC methods that we also applied to the real data. See Supporting Information 3 for more details.

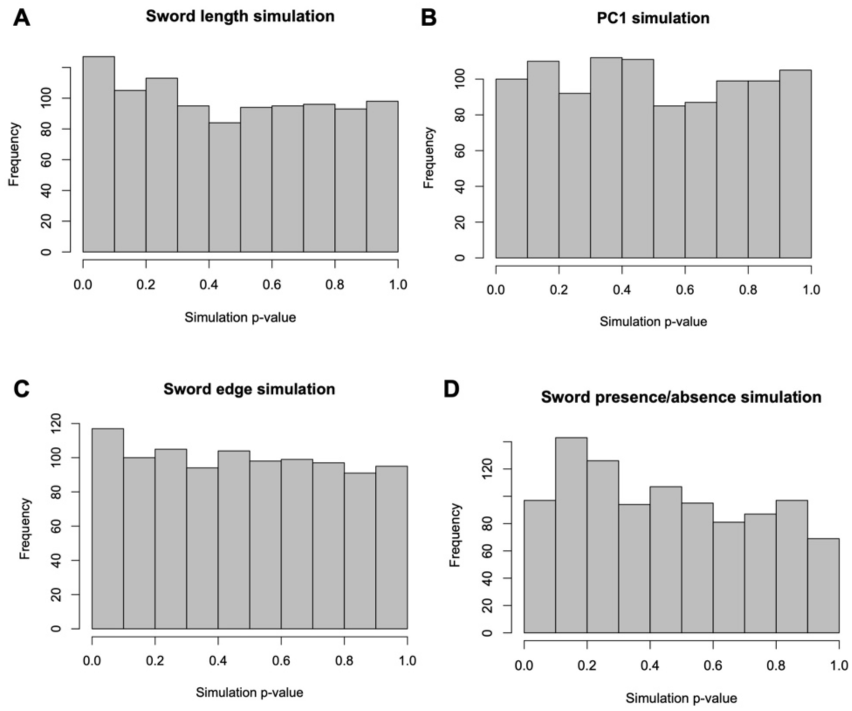


**Fig. S12.** P-value distribution from PIC analysis of 1,000 simulations of continuous (A-C) and binary (D) traits using the *Xiphophorus* phylogeny. For each simulation, we evaluated the correlation between observed body size and the simulated trait and recorded the p-value after phylogenetic correction. Simulations were performed such that the mean and variance of the simulated traits matched the observed traits in our dataset. We find some evidence for p-value inflation in these distributions for simulations of continuous traits, but it is mild. In **A**) 6% of simulations had a p-value less than 0.05, in **B**) 5.5% of simulations had a p-value less than 0.05, and in **C**) 6.5% of simulations had a p-value less than 0.05. For our simulations of binary traits shown in **D,** (intended to match the distribution of sword presence or absence), the p-value distribution was somewhat conservative and only 2% of simulations had a p-value less than 0.05. See Supporting Information 3 for more details.

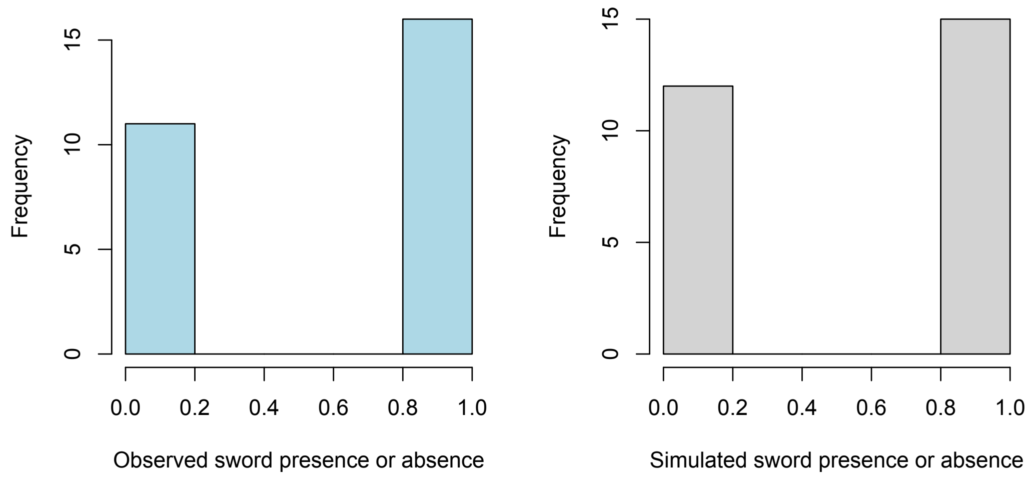


**Fig. S13.** Comparison of observed distribution of sword presence and absence in the *Xiphophorus* phylogeny to simulations of sword presence and absence. We used these simulations to evaluate the performance of phylogenetic logistic regression methods applied to the real data. See Supporting Information 3 for more details.

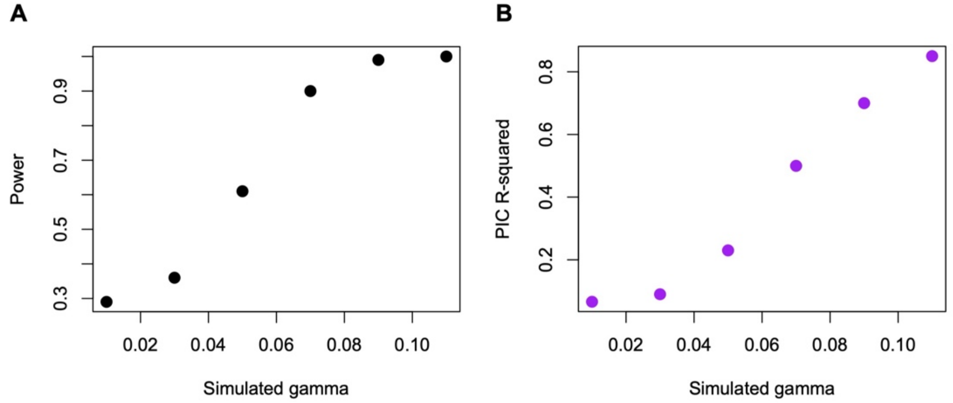


**Fig. S14.** Power simulation results for correlated evolution of simulated traits. We varied the gamma parameter in modeling trait coevolution from 0.01-0.11 and performed 1,000 simulations of two coevolving traits on the swordtail phylogeny for each gamma value. **A)** For each pair of simulated traits, we applied PIC correction as we had for the real data, and measured in what proportion of simulations traits were correlated at $p<0.05$. This proportion is plotted as power on the y-axis. **B**) For the same set of simulations, we plot the median $R^2$ inferred in PIC analysis for those 1,000 simulations. In the real data, the median significant correlation we detect is $R^2$ of 0.54. This hints that we have modest power in our analysis of some traits in our dataset.

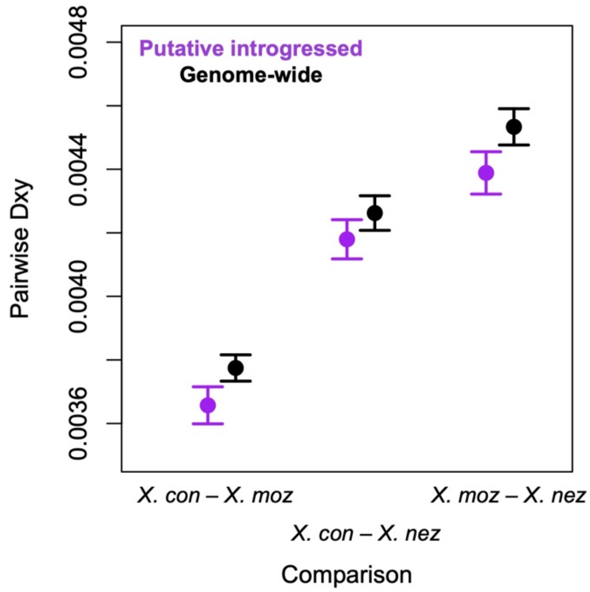


**Fig. S15**. Pairwise divergence between *X. continens*, *X. montezumae*, and *X. nezahualcoyotl* genome-wide (black) and in regions identified as putatively introgressed based on PhyloNetHMM analysis (purple). Whiskers represent two standard errors. See Supplementary Information 4 for more information.

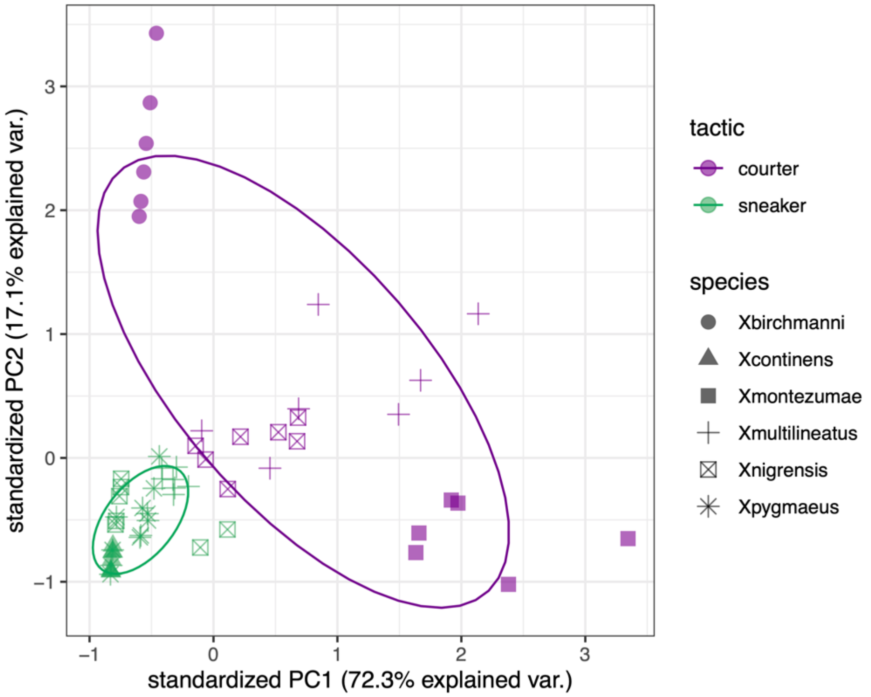


**Fig. S16.** PCA analysis of male phenotypes for a subset of Northern swordtail fish species. This figure is similar to that presented in Figure 2B in the main text except that it includes *X. birchmanni*, a species that has lost the sword but that has a number of other male ornaments as well as large body size. Color indicates mating strategy and shape indicates species. Ellipses indicate samples that falls within ± 1 standard deviation of the mean of that group.

### Supporting Information References

Cui, R., Schumer, M., Kruesi, K., Walter, R., Andolfatto, P. & Rosenthal, G. (2013) Phylogenomics reveals extensive reticulate evolution in Xiphophorus fishes. *Evolution*, **67**, 2166–2179.

Díaz-Uriarte, R. & Garland, T., Jr. (1996) Testing Hypotheses of Correlated Evolution Using Phylogenetically Independent Contrasts: Sensitivity to Deviations from Brownian Motion. *Systematic Biology*, **45**, 27–47.

Garland, T., Jr., Harvey, P.H. & Ives, A.R. (1992) Procedures for the Analysis of Comparative Data Using Phylogenetically Independent Contrasts. *Systematic Biology*, **41**, 18–32.

Ives, A.R. & Garland, T., Jr. (2010) Phylogenetic Logistic Regression for Binary Dependent Variables. *Systematic Biology*, **59**, 9–26.

Langdon, Q.K., Powell, D.L., Kim, B., Banerjee, S.M., Payne, C., Dodge, T.O., *et al.* (2022) Predictability and parallelism in the contemporary evolution of hybrid genomes. *PLOS Genetics*, **18**, e1009914.

Li, H. & Durbin, R. (2011) Inference of human population history from individual whole-genome sequences. *Nature*, **475**, 493–496.

Liu, K.J., Dai, J., Truong, K., Song, Y., Kohn, M.H. & Nakhleh, L. (2014) An HMM-Based Comparative Genomic Framework for Detecting Introgression in Eukaryotes. *PLOS Computational Biology*, **10**, e1003649.

Liu, L., Yu, L., Kubatko, L., Pearl, D.K. & Edwards, S.V. (2009) Coalescent methods for estimating phylogenetic trees. *Molecular Phylogenetics and Evolution*, **53**, 320–328.

Paradis, E. & Schliep, K. (2019) ape 5.0: an environment for modern phylogenetics and evolutionary analyses in R. *Bioinformatics*, **35**, 526–528.

Powell, D.L., Moran, B.M., Kim, B.Y., Banerjee, S.M., Aguillon, S.M., Fascinetto-Zago, P., *et al.* (2021) Two new hybrid populations expand the swordtail hybridization model system. *Evolution*, **75**, 2524–2539.

Revell, L.J. (2012) phytools: an R package for phylogenetic comparative biology (and other things). *Methods in Ecology and Evolution*, **3**, 217–223.

Schumer, M., Cui, R., Powell, D.L., Rosenthal, G.G. & Andolfatto, P. (2016) Ancient hybridization and genomic stabilization in a swordtail fish. *Molecular Ecology*, **25**, 2661–2679.

Schumer, M., Xu, C., Powell, D.L., Durvasula, A., Skov, L., Holland, C., *et al.* (2018) Natural selection interacts with recombination to shape the evolution of hybrid genomes. *Science*, **360**, 656.

Tung Ho, L. si & Ané, C. (2014) A Linear-Time Algorithm for Gaussian and Non-Gaussian Trait Evolution Models. *Systematic Biology*, **63**, 397–408.
